# A tumour-promoting senescent secretome triggered by platinum chemotherapy exploits a targetable TGFβR1/Akt-mTOR axis in lung cancer

**DOI:** 10.1101/2022.08.01.502019

**Authors:** Estela González-Gualda, David Macias, Samir Morsli, José Ezequiel Martín, Hui-Ling Ou, Mary Denholm, Ioana Olan, Reuben Hoffmann, Mark Dane, Dimitris Veroutis, Guillermo Medrano, Francisca Mulero, Carla P. Martins, Mariano Barbacid, Vassilis Gorgoulis, James E. Korkola, Doris M. Rassl, Gary J. Doherty, Robert C. Rintoul, Masashi Narita, Daniel Muñoz-Espín

## Abstract

Platinum-based chemotherapy is commonly used for non-small cell lung cancer (NSCLC) treatment, yet clinical outcomes remain poor. Cellular senescence and its associated secretory phenotype (SASP) can have multiple tumour-promoting activities, although these are largely unexplored in lung cancer. Here we show that cisplatin-derived SASP enhances the malignant phenotype of lung cancer cells. Using xenograft, orthotopic and Kras^G12V^-driven murine NSCLC models, we demonstrate that cisplatin-induced senescent cells strongly promote tumour progression. Mechanistically, we find that a TGF-β-enriched SASP drives pro-proliferative effects through TGFβR1 and Akt/mTOR pathway activation. We validate the translational relevance of chemotherapy-induced SASP using clinical NSCLC samples from patients who received neoadjuvant platinum-based chemotherapy. Importantly, TGFβR1 inhibition with galunisertib or senolytic treatment significantly reduces tumour promotion driven by cisplatin-induced senescence. Finally, we demonstrate, using distinct murine NSCLC models, that addition of TGFBR1 inhibitors to platinum-based chemotherapy reduces tumour burden and improves survival, providing pre-clinical proof-of-concept for future trial designs.

## Introduction

Lung cancer accounts for the highest proportion of cancer-related deaths worldwide (over 1.6 million deaths annually). Non-Small Cell Lung Cancer (NSCLC) accounts for around 85% of all lung cancer diagnoses, with adenocarcinoma being the most common histological subtype. Significant advances over the last two decades have been made towards improving lung cancer treatment. However, the overall survival and cure rates for NSCLC remain low, with treatment resistance and progression being unavoidable in most of cases, particularly in patients with advanced disease^1^. Although the use of novel molecularly targeted therapies and immunotherapy has led to significantly improved outcomes in selected patients, platinum-based doublet therapy (cisplatin or carboplatin in combination with another cytotoxic drug), either alone or in combination with anti-PD-(L1) immunotherapy, remains standard treatment (including first-line treatment) in the neoadjuvant, adjuvant, locally advanced and metastatic disease settings^1^. Despite favourable clinical activity, most patients harbour at least minimal residual disease and progress or relapse shortly after treatment completion, with 75% of lung cancer patients dying within 5 years from first diagnosis^1^. Therefore, continued research into the mechanisms underlying treatment response and failure is crucial to develop enhanced therapeutic regimens to improve NSCLC patient outcomes.

Early studies showed that platinum-based chemotherapy results in increased levels of intratumoral Senescence Associated β-galactosidase (SA-β-gal) activity in NSCLC patient tissue samples, a marker commonly associated with the onset of cellular senescence^2^. Additional studies have further demonstrated the accumulation of senescent cells in different human tumour types after neoadjuvant chemotherapy, including breast cancer, mesothelioma, prostate cancer, renal cell carcinoma and rectal cancer^3^, suggesting that it is a common response to anti-cancer therapies. Senescence is a cell programme elicited in response to a variety of stimuli, including DNA damage, oncogene activation, and therapy-induced genotoxic stress, whereby proliferating cells become stably arrested and undergo a series of structural and metabolic changes that impacts tissue homeostasis^4, 5^. A key hallmark of this cellular state, affecting both cancer and non-malignant cells, is the implementation of a strong paracrine secretion of cytokines, growth factors, proteases and other mediators, jointly named the Senescence-Associated Secretory Phenotype (SASP), which creates a local inflammatory milieu that affects the surrounding tissue^6, 7^ Senescence is considered a tumour suppressive mechanism as it prevents the propagation of damaged cells, but increasing evidence indicates that when senescent cells persist in tissues they can also drive potent tumour-promoting effects, including increased proliferation of premalignant and cancer cells, invasion, angiogenesis, metastasis and even immunosuppression in a paracrine manner^8, 9^. Recent investigations have primarily focused on the mechanisms by which cellular senescence can contribute to cancer development and the side effects of cancer therapies^10, 11^, but their impact in the particular context of the lung remains largely unknown.

Despite a general list of factors commonly proposed to constitute the SASP, including IL-6, IL-1α, CXCL2, MMP-3 and VEGF, among many others, its composition is known to be very heterogeneous and dynamic, and hence highly dependent on the inducer, the tissue of origin or cell type and even the duration of the senescent burden^8^. Transforming Growth Factor-β (TGF-β) is a pleiotropic cytokine that governs a wide array of cellular mechanisms in physiological and pathological processes, and it has also been reported to be secreted by senescent cells, contributing to paracrine-induced senescence, fibrosis and immunomodulation^12–14^. In the context of cancer, TGF-β has been described to play a dual role, the result of which largely depends on the surrounding microenvironment and the recipient cell. Paradoxically, this cytokine functions as a tumour suppressor in pre-malignant cells by inducing cell cycle arrest and apoptosis, but it can also drive tumorigenic and pro-metastatic responses in malignant cells through the promotion of Epithelial-to-Mesenchymal Transitions (EMT), tumour invasion, proliferation and immune suppression^15^. The complexity of the contextual nature of TGF-β as part of the SASP and its potential connection with chemotherapy-induced senescence remains unexplored and warrants further investigation.

In this study, we demonstrate that cisplatin treatment results in the induction and accumulation of senescent cells in human and murine lung adenocarcinomas. Comprehensive phenotypic assessment reveals that the SASP derived from human and murine cisplatin-induced senescent lung cancer cells drives the acquisition of tumour-promoting traits in a paracrine manner. This was then recapitulated in lung cancer human xenograft, orthotopic and Kras^G12V^-driven mouse models. Mechanistically, we show that exposure of lung cancer cells to cisplatin-derived SASP orchestrates the TGFβR1-driven activation of Akt/mTOR signalling, which is responsible for the induction of increased proliferation and malignant traits. Of translational relevance, we provide mechanistic validation by histological analyses of tissue sections from NSCLC patients subjected to neoadjuvant platinum-based therapy. Moreover, we determine that pharmacological inhibition of TGFβR1 with galunisertib treatment reverts the tumour promoting effects of cisplatin-induced senescence both *in vitro* and *in vivo*. Finally, we propose a novel therapeutic modality combining chemotherapeutic cisplatin treatment together with TGFβR1 inhibition, which demonstrates improved treatment outcomes, reduced tumour burden and enhanced survival in different lung tumour-bearing mouse models.

## Results

### Cisplatin-induced senescence promotes malignant traits in recipient cancer cells through the secretion of SASP factors

Mounting evidence has shown that therapy-induced senescence can drive a chronic inflammatory niche in the tumour microenvironment, which in turn has the potential to promote a variety of tumour-promoting activities^8, 16^. To investigate the effects of chemotherapy-derived SASP in the context of NSCLC, human lung adenocarcinoma A549 cells were subjected to cisplatin (15 μM), docetaxel (200 nM) and palbociclib (15 μM) treatment. The analysis of increased SA-β-gal staining, stable proliferation arrest and the expression markers of senescence at RNA and protein levels confirmed the implementation of senescence upon drug exposure (Fig. 1a,b; Extended Data Fig. 1a,b) . First, medium was conditioned for 48 h with each type of therapy-induced senescent cells, and untreated A549 cells were exposed to each type of conditioned medium (CM) (Fig. 1c). Our analyses revealed that exposure to cisplatin-derived SASP resulted in increased cell proliferation and cell migration, compared to control CM and docetaxel-and palbociclib-derived SASPs, with exposed cells achieving an average 85.4% confluency at 42 h in the presence of cisplatin-induced senescent CM versus 53.2% in control conditions (Fig. 1d, left). Cell tracking during a wound-healing scratch assay showed that exposure to cisplatin-SASP also significantly increased A549 cell migration, compared to all other conditions (Fig. 1d, right). To exclude the potential effect of acidification or lower nutrient levels in CMs, we analysed the pH and glucose concentration and no changes were observed (Extended Data Fig. 1k,l). We next showed that exposure to cisplatin-induced senescent CM for 10 days led to a 3.4-fold increase in the number of colonies formed compared to control CM, while palbociclib-induced senescent CM had no effect and docetaxel-derived SASP resulted in significantly fewer colonies (Fig. 1e). The number of spheres formed in low-attachment conditions was also significantly higher in the presence of cisplatin-derived SASP compared to all other conditions (Fig. 1f). Notably, these spheres also exhibited irregular shapes as well as the formation of protrusions, which, together with the enhanced migration properties observed in our wound-healing analyses, suggested the acquisition of an invasive phenotype consistent with an Epithelial-to-Mesenchymal Transition (EMT). To determine the effect of the direct interaction between senescent and untreated tumour cells, we cultured A549 cells in a three-dimensional (3D) spheroid system alone or with chemotherapy-induced senescent cells. Notably, 3D co-culture with cisplatin-induced senescent A549 cells resulted in the development of significantly larger spheres compared to all other conditions (Fig. 1g). To better understand the changes occurring in cells upon exposure to cisplatin-derived SASP, we subjected A549 cells to a mitochondrial stress metabolic assay after incubation with CM for 48 h. Oxygen Consumption Rate (OCR) analysis upon injection of drugs targeting different complexes of the mitochondrial respiratory chain revealed a significantly increased maximal respiration in cells exposed to cisplatin-derived SASP compared to control (Fig. 1h), suggesting a metabolic rewiring characterised by an increased ability to appropriately respond to high demands of ATP, and therefore a higher endurance during stress periods. No changes in respiratory ability were detected after the incubation of A549 cells with docetaxel- or palbociclib-induced senescent CM (Extended Data Fig. 1n). Interestingly, bioenergetic profiling of control and chemotherapy-induced senescent A549 cells revealed that senescent cells also harbour an increased maximal respiratory capacity compared to control cells, with cisplatin-induced senescent cells showing the highest OCR levels among the three anti-cancer therapies (Extended Data Fig. 1m, left).

**Figure 1.**
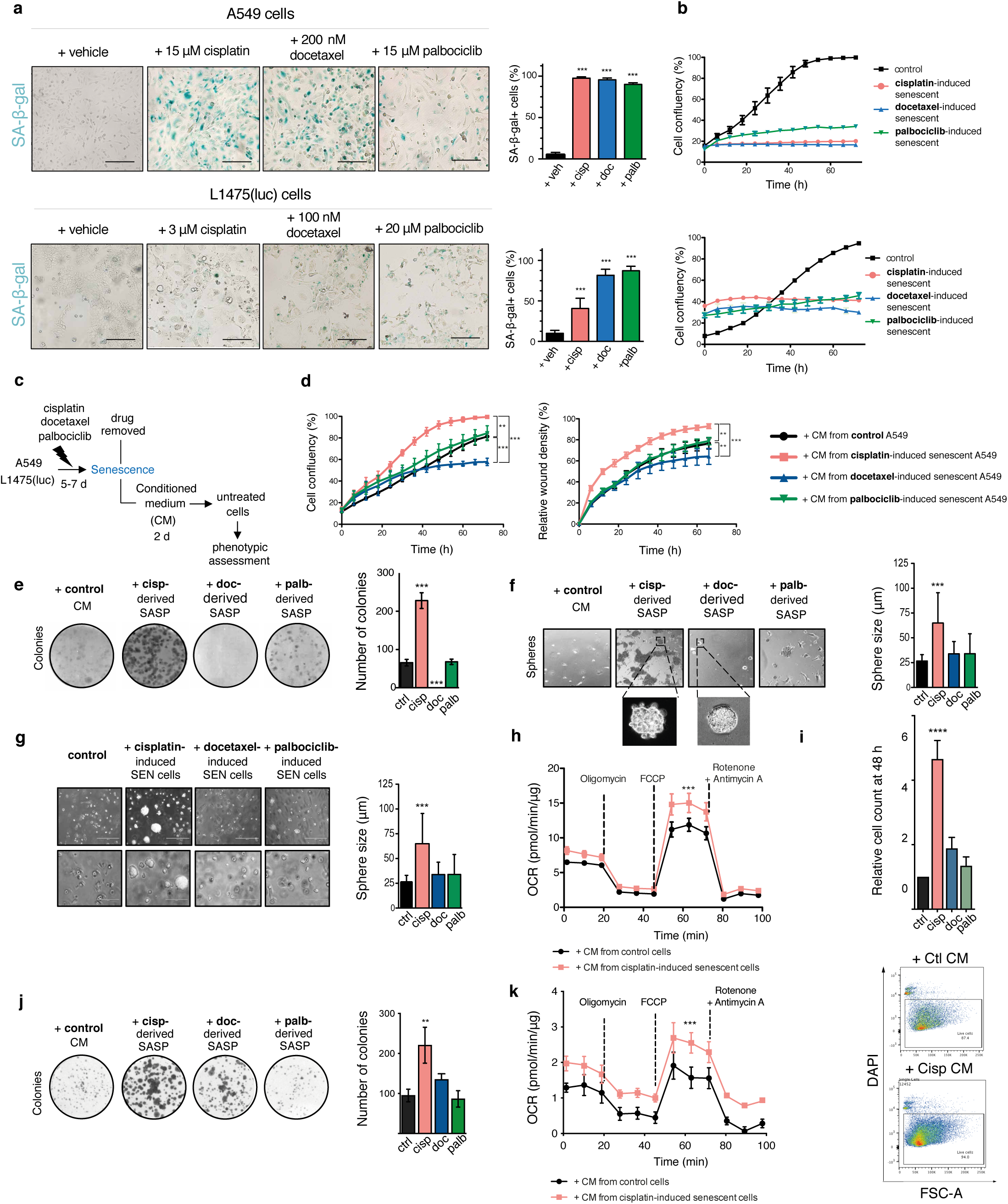
Cisplatin-induced senescence promotes malignant traits on non-senescent cancer cells through the secretion of SASP factors. **a.** Representative images of control and chemotherapy-treated A549 cells or L1475(luc) cells fixed and stained for SA-β-gal activity after 7 days of treatment. Scale bar = 100 µm. Right, proportion of SA-β-gal+ A549 cells or L1475(luc) cells for each treatment. **b.** Cell confluency over time of control and senescent A549 cells or L1475(luc) cells after 7 days of treatment, respectively. **c.** Schematic representation of experimental layout to test the effect of chemotherapy-derived SASP on non-senescent cells. A549 cells were treated with 15 µM CDDP, 150 nM docetaxel or 15 µM Palbociclib for 7 days for senescence induction. Senescent cells were subsequently washed and fresh medium was added and conditioned for 48 h. Conditioned Media (CM) were collected, centrifuged and added onto non-senescent A549 cells, which were subsequently assayed for phenotypic assessment. **d.** Left, A549 cell confluency (%) over a 72h incubation-period exposed to CM from control or chemotherapy-induced senescent cells. Right, relative wound density of migrating A549 cells after scratching over a 66h period exposed to CM from control or chemotherapy-induced senescent cells. **e.** Left, representative images of A549 colonies formed after 10 days upon the exposure to the different CMs from non-senescent and chemotherapy-induced senescent cells. Right, quantification of the number of colonies for each experimental condition. **f.** Left, representative images of A549 3D tumour spheres formed after 7 days of two-phase co-culture with control or chemotherapy-induced senescent cells. Right, quantification of sphere size in each experimental condition. **g.** Left, representative images of A549 spheres formed after 10 days upon the exposure to the different CMs from non-senescent and chemotherapy-induced senescent cells. Right, quantification of the number of spheres for each experimental condition **h.** Oxygen Consumption Rate (OCR) of non-senescent A549 cells at basal conditions and upon injection of oligomycin, FCCP, rotenone and antimycin A, measured after 72 h incubation with CM from control or cisplatin-induced senescent A549 cells. **i.** L1475(luc) cell count relative to control condition of cells exposed to control- or chemotherapy-induced senescent CM for 48. Bottom, representative histograms depicting live cell events of samples upon exposure to control- or cisplatin-induced senescent CM. **j.** Left, representative images of L1475(luc) colonies formed upon exposure to control- or chemotherapy-induced senescent CM for 10 days. Right, quantification of the number of colonies formed in each experimental condition. **k**. OCR of non-senescent murine L1475(luc) cells at basal conditions and upon injection of oligomycin, FCCP, rotenone and antimycin A after 72 h incubation with CM from control or cisplatin-induced senescent L1475(luc) cells. All data are plotted as mean ± SD (n = 3), except for OCR plots in **h** and **k**, where data are shown as mean ± SD from one representative experiment (out of three independent experiments performed). Statistical significance was assessed using two-tailed one-way or two-way ANOVA, followed by Tukey’s multiple comparisons test, **p < 0.01, ***p < 0.005, ****p < 0.001.

We next aimed to validate our findings using the primary KRas^G12D/WT^;p53^-/-^ (KP) murine L1475(luc) cell line, induced to cellular senescence by cisplatin (3 μM), docetaxel (100 nM) or palbociclib (20 μM) treatments (Fig. 1a,b; Extended Data Fig. 1c,d). Relative cell count upon 48 h of exposure to CMs demonstrated that cisplatin-induced senescent CM also increased proliferation of untreated L1475(luc) cells compared to all other conditions (Fig. 1i). In addition, culture of L1475(luc) with cisplatin-induced senescent CM for 10 days resulted in a higher number of colonies formed (Fig. 1j), and respiratory profiling of cells exposed to this CM revealed a significantly enhanced maximal respiration (Fig. 1k), in line with our findings in A549 cells (Fig. 1h). In contrast with A549 cells, L1475(luc) cells exposed to docetaxel- and palbociclib-induced senescent CM also showed increased OCR levels, albeit to a lower extent compared with cisplatin-derived SASP exposure (Extended Data Fig. 1o). Bioenergetic analysis of control and senescent L1475(luc) cells showed trends similar to those observed in A549 cells, with the exception of docetaxel-induced senescent cells, which presented a respiratory profile comparable to control cells (Extended Data Fig. 1m, right).

In order to explore whether the cisplatin analogue carboplatin also resulted in similar phenotypes, we first determined the induction of senescence in A549 cells upon 7 day treatment with 7.5 μM carboplatin (Extended Data Fig. 1e,f). When untreated A549 cells were exposed to carboplatin-derived SASP, colony-formation analyses demonstrated a significant stimulation of the formation of colonies compared to control CM (Extended Data Fig. 1,g). Finally, to further validate our observations in an alternative model where platinum-based treatment is standard-of-care, we subjected ovarian cancer PEO4 cells to cisplatin and carboplatin treatments (2.5 μM and 10 μM, respectively) and validated the induction of cellular senescence (Extended Data Fig. 1h,i). Importantly, untreated PEO4 cells also exhibited increased proliferation rates upon the exposure to both cisplatin- and carboplatin-derived SASPs, versus control (Extended Data Fig. 1j).

Altogether, our *in vitro* assessments with both human and murine lung adenocarcinoma cells show that platinum-based treatment results in a particular senescence subtype that can exacerbate malignant properties in recipient, previously untreated, lung cancer cells through the SASP. This promoted tumour cell proliferation, migration, sphere formation, as well as enhanced respiratory endurance.

### Cisplatin-induced senescent A549 and L1475(luc) cells support increased tumour growth in xenograft and orthotopic models of NSCLC

To validate the ability of cisplatin-induced senescence to drive increased tumour cell proliferation in physiological models, we first subcutaneously co-transplanted cisplatin-induced senescent A549-GFP+ cells together with untreated A549-mCherry+ cells and analysed the growth of the xenografts formed over time (Fig. 2a). As shown in Fig. 2b, relative tumour volume was significantly higher when tumour cells were co-transplanted with senescent cells, as compared to untreated cells transplanted alone. Strikingly, by day 21, co-transplanted tumours (untreated + senescent cells) were on average 2.18 times the volume of control tumours (average tumour volume of co-transplanted: 325.44 +/- 116.82 mm^3^; control: 149.00 +/- 60.44 mm^3^; N=18 tumours/group) (Fig. 2b). Transplantation of senescent cells alone resulted in xenograft recession, confirming that the effect in co-transplanted tumours is likely driven through paracrine support from senescent cells and not because of senescence escape. Mean tumour weight of co-transplanted xenografts (251.11 +/- 87.30 mg, N=18) was also significantly higher than untreated tumours (156.67 +/-78.74 mg, N=18) and senescent tumours (37.22 +/- 17.08 mg, N=18) (Extended Data Fig. 2a). Histological analyses of the tumours confirmed decreased levels of the proliferative marker Ki67 and increased expression of p21 in senescent xenografts compared to untreated and co-transplanted tumours (Extended Data Fig. 2b). Representative pictures of the resected tumours at experiment completion are shown in Fig. 2c.

**Figure 2.**
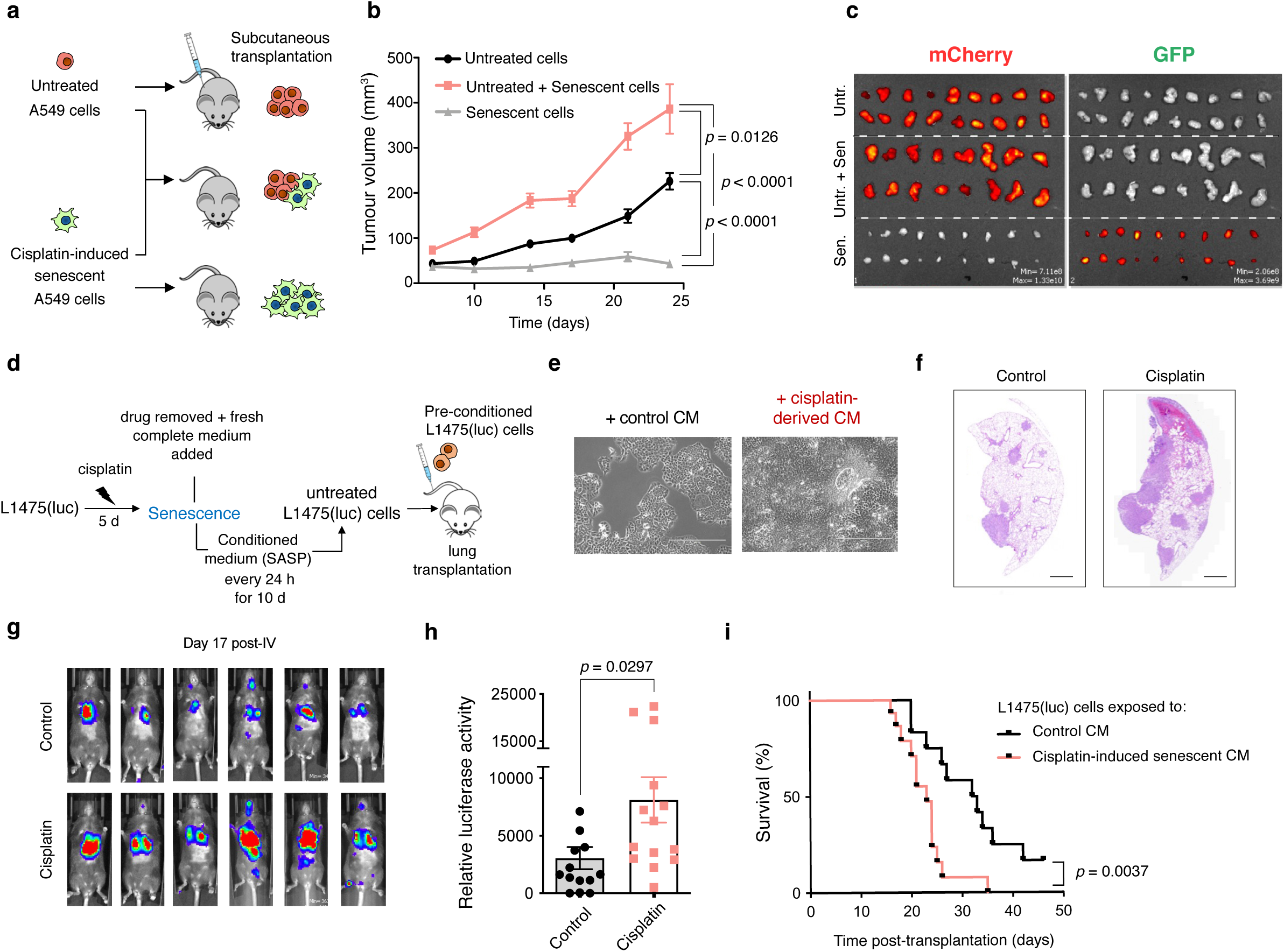
Cisplatin-induced senescent A549 and L1475(luc) cells support increased tumour growth in xenograft and orthotopic models of NSCLC. **a.** Schematic representation of experimental layout using the subcutaneous xenograft model. Briefly, animals were transplanted with either 4x10^6^ untreated A549-mCherry+ cells, 4x10^6^ untreated A549-mCherry+ cells and 1x10^6^ cisplatin-induced senescent A549-GFP+ cells, or 4x10^6^ cisplatin-induced senescent A549-GFP+ cells alone in the flank. **b.** Tumour volume of xenografts over time. Data is shown as mean ± SEM (n=18 tumours per group). **c.** Fluorescent images of resected tumours at the end of the experiment. **d.** Schematic representation of experimental layout. Briefly, untreated L1475(luc) cells were continuously exposed to the CM from control- or cisplatin-induced senescent cells for 10 days prior to lung transplantation via tail-vein injection. **e.** Representative images of L1475(luc) cells exposed to control- and cisplatin-induced senescent CM prior to transplantation. Scale bar = 400 µm. **f.** Representative images of histological sections from lungs resected at 17 days post-transplantation of lung cancer cells in each experimental condition. Scale bar = 1 mm. **g.** Representative images of luciferase activity in mice 17 days after transplantation with lung cancer cells exposed to control- or cisplatin-induced senescent cell CM. **h.** Quantification of luciferase activity at day 17 post-transplantation, relative to activity recorded on day 1 after cell transplantation (n=14 per group). **i.** Survival curve of mice in each experimental group (n=14 per group). Data in **b** and **h** are shown as mean ± SEM. Statistical significance for tumour volume was assessed using two-way ANOVA, followed by Tukey’s multiple comparisons test, and by two-tailed Student’s t-test for relative luciferase activity. Survival analysis was performed using the Kaplan-Meier method and a two-sided log-rank test was conducted to determine statistical significance. Orthotopic model data represents three independent experiments.

We next investigated the paracrine effects of cisplatin-induced senescence using a KRas^G12D/WT^;p53^-/-^ orthotopic murine model. L1475(luc) cells were exposed to control or cisplatin-induced senescent CM for 10 days and were subsequently transplanted in the lungs of C57BL/6 mice via tail-vein injection (Fig. 2d). The expression of luciferase in these cells allowed the analysis of tumour burden over time via D-luciferin administration and bioluminescence imaging. As an internal control, cells exposed to cisplatin-derived SASP proliferated faster than those exposed to control CM in culture (Fig. 2e; Extended Data Fig. 2c). We observed that the transplantation of cells that had been exposed to cisplatin-derived SASP resulted in a significantly greater luciferase activity in the lungs, relative to day 1 post-injection (Fig. 2h). Representative images of luciferase signal at 17 days post-transplantation are shown in Fig. 2g, and a higher number and greater size of tumour foci in lungs transplanted with cells exposed to cisplatin-derived SASP at the same time-point can be observed in Fig. 2f (stained with haematoxylin and eosin). Notably, the transplantation of tumours exposed to cisplatin-induced senescent CM resulted in significant decrease in median survival by 30% (23 days compared to 32.5 in mice transplanted with tumours conditioned with control CM; Fig. 2i).

Chemotherapy is known to be gerontogenic and can induce senescence affecting both tumour and non-cancer tissues^17^, while senescence has been shown to promote adverse effects of chemotherapy and cancer relapse^18^. Considering the well-known increase of senescence burden in tissues during ageing and upon genotoxic stress we decided to investigate the potential interplay between cisplatin-induced senescence, the process of ageing and lung tumour progression^19^. Toward this aim, young and middle-aged mice were subjected to two cycles of cisplatin treatment and then L1475(luc) cells were transplanted orthotopically in the lungs (Extended Data Fig. 2d). Subsequent RT-qPCR analysis of RNA extracted from whole lungs at day 14 post-treatment revealed evidence of increased gene expression levels of the senescence markers *Cdkn1a*, *Il6* and *Tgfb2* in cisplatin-treated animals compared to the vehicle group (Extended Data Fig. 2e). This observation correlates with our previous work showing that cisplatin treatment post-orthotopic transplantation of L1475(luc) cancer cells also results in the onset of senescence in the lung^20^. Tumour burden analysis by bioluminescence imaging revealed significantly higher relative luciferase activity 14 days after tumour transplantation in middle-aged animals compared to young individuals (Extended Data Fig. 2f). Representative images of luciferase signal at day 7 and 14 post-transplantation of cells are shown in Extended Data Fig. 2g. Of note, median survival in aged animals was decreased by 31% (14.5 days compared to 21 days in young individuals; Extended Data Fig. 2h). Finally, in order to gain further insights into the impact of platinum-based therapy, we analysed body weight over time as an indirect measure of chemotherapy-related adverse effects and observed no major differences between the groups during cisplatin administration. In contrast, aged individuals suffered a marked weight loss shortly after tumour transplantation, while young mice maintained more stable relative weight during the same time frame (Extended Data Fig. 2i). These results suggest that young individuals present higher tolerability towards cisplatin treatment, and that ageing exacerbates tumour progression.

Taken together, our findings confirm that the induction of senescence in response to cisplatin treatment in mice exerts detrimental effects in a paracrine fashion, boosting lung cancer progression and shortening lifespan.

### Transcriptomic and proteomic analyses reveal chemotherapy-context dependent differences in SASP signatures and that TGF-β ligands are overrepresented in cisplatin-derived SASP

To gain insight into the gene expression profiles occurring upon chemotherapy-driven senescence, we next performed RNA-seq analyses of control, cisplatin-, docetaxel- and palbociclib-induced senescent A549 cells. Our analyses showed increased expression of known senescence mediators such as *TP53* (p53) and *CDKN1A* (p21^WAF1/Cip1^) genes, while reduced expression of genes promoting cell cycle progression (such as *E2F*, *CDC* and *CDK* genes) and DNA replication (such as *PCNA*, *POL* and *MCM* genes), which correlates with the implementation of a stable cell cycle arrest (see scaled expression profile in Fig. 3a; Extended Data Fig. 3a). Gene-set enrichment analysis (GSEA) furthered these observations by revealing that cisplatin-, docetaxel- and palbociclib-treated A549 cells are all positively enriched for signatures of senescence and negatively enriched for cell cycle and DNA replication pathways that promote proliferation compared to vehicle-treated cells, thereby confirming the implementation of the senescent programme (Extended Data Fig. 3a). As expected, cisplatin-treated cells displayed much greater transcription of mismatch repair and DNA repair-related genes. Notably, palbociclib-induced senescent cells presented the most distinct transcriptional profile compared to cisplatin- and docetaxel-induced senescent A549 cells. We next sought to uncover the genes exclusively upregulated in each type of chemotherapy-driven senescence (Fig. 3b), and observed that docetaxel-induced senescent cells present the highest number of uniquely upregulated genes (1961), versus 751 genes in cisplatin-induced senescence and 363 genes in palbociclib-induced senescent cells.

**Figure 3.**
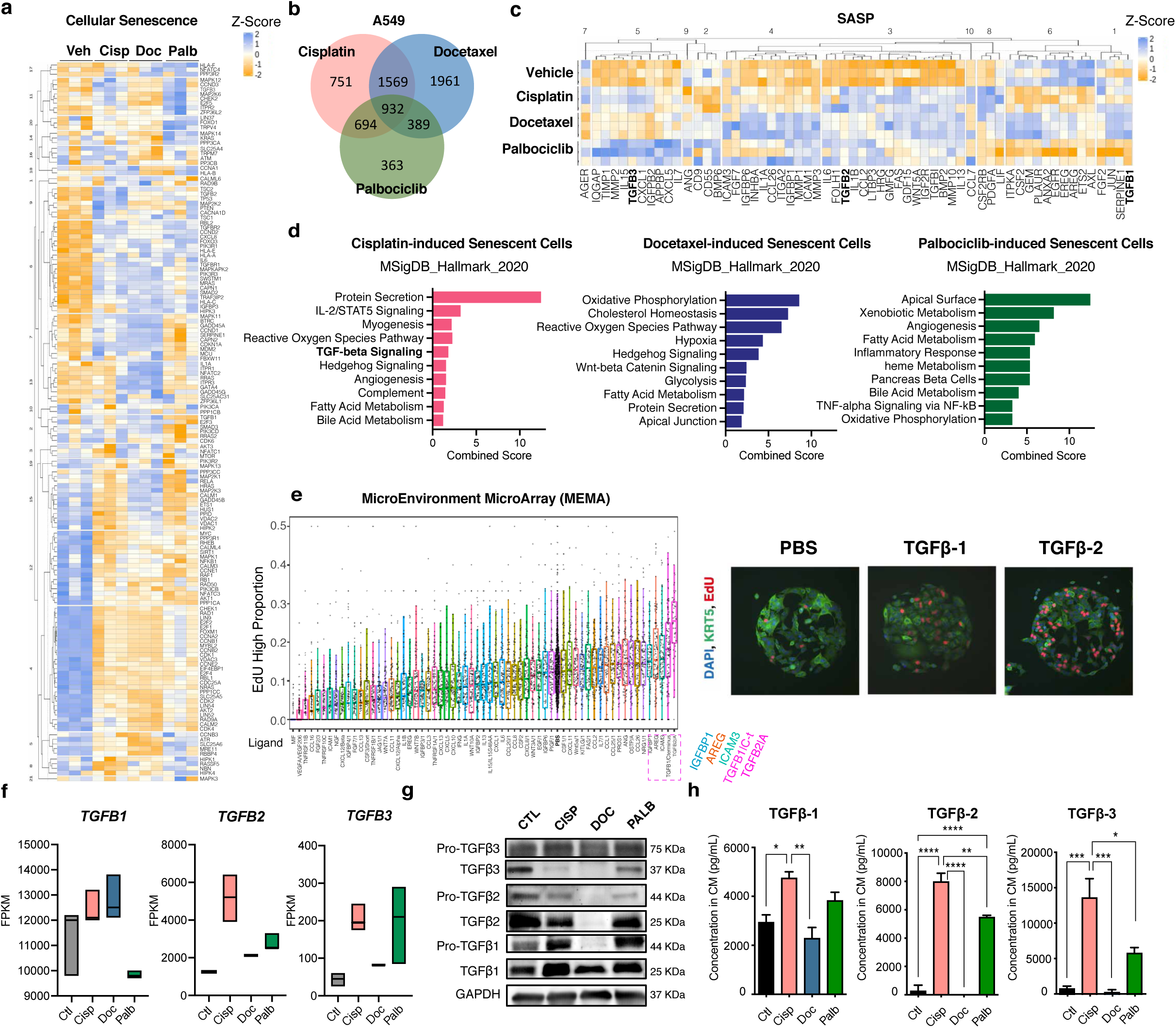
Transcriptomic and proteomics analyses reveal different transcriptional and SASP signatures of chemotherapy-induced senescent cells and highlight TGFβ ligands as potential candidate drivers of malignant traits in cisplatin-derived SASP. **a.** Heatmap displaying expression z-scores of most significantly altered gene expression changes in cellular senescence pathway (hsa04218) and their hierarchical clustering in control and cisplatin-, docetaxel- and palbociclib-induced senescent A549 cells. **b.** Venn diagram of the number of genes significantly upregulated in cisplatin-, docetaxel- and palbociclib-induced senescent A549 cells vs control. **c.** Heatmap of expression z-scores of selected SASP genes in control and cisplatin-, docetaxel- and palbociclib-induced senescent A549 cells. **d.** Top differentially enriched pathways in MSigDB Hallmark 2020 collection, sorted by combined score ranking, in cisplatin-, docetaxel- and palbociclib-induced A549 senescent cells vs control. **e.** Left, proportion of EdU high intensity A549 cells upon 72 h of exposure to different SASP ligands using the MEMA technology, ranked left to right form lowest induced proportion to highest. Right, representative immunofluorescence images of MEMA wells containing A549 cells upon exposure to PBS, TGFβ-1 and TGFβ- 2 recombinant ligands, depicting DAPI and KRT5 staining and EdU incorporation. **f.** Fragments per Kilobase of transcript per Million mapped reads (FPKM) of *TGFB1*, *TGFB2* and *TGFB3* ligands in control and cisplatin-, docetaxel- and palbociclib-induced senescent A549 cells. **g.** Western blot analysis of whole cell extracts of control and cisplatin-, docetaxel- and palbociclib-induced senescent A549 cells. **h.** Average concentration in CM of TGFβ1, TGFβ2 and TGFβ3 free ligands (activated) in each experimental condition measured by ELISA. Data are shown as mean ± SD (n=3). Statistical significance was assessed by two-tailed one-way ANOVA, followed by Tukey’s multiple comparisons test, *p < 0.05, **p < 0.01, ***p < 0.005, ****p < 0.001.

To dissect gene expression profile changes in the secretory phenotype of each type of senescence, we next investigated the expression of known SASP factors in vehicle- and chemotherapy-treated A549 cells (Fig. 3c). As expected, we observed a general increase in the transcription of inflammatory ligands and proteases in senescent A549 cells in most of the clusters, such as genes encoding for IL6, IL1B, CCL2 and MMPs. Clusters 3 and 8 appeared particularly enriched in cisplatin-induced senescent cells versus docetaxel- and palbociclib-induced senescent and control A549 cells, which included genes encoding for TGFB2, CSF2RB, PDGFA and LIF. To further understand the most significant changes at the signalling level upon senescence induction in A549 cells, we performed pathway enrichment analyses of significantly differentially expressed genes in each type of chemotherapy-induced senescent cells against pre-defined sets from the Molecular signatures database (MSigDB). This revealed a significant enrichment in metabolic pathways across the three types of senescence (Fig. 3d). Interestingly, biologically relevant pathways including IL-2/STAT5 and TGF-β pathways were found to be uniquely enriched in cisplatin-induced senescent cells.

Transcriptomic analysis of murine L1475 lung adenocarcinoma cells upon chemotherapeutic treatment revealed that cisplatin- and docetaxel-induced senescent cells present marked distinct gene expression profiles compared to palbociclib- and vehicle-treated cells (Extended Data Fig. 3c,d). Similarly to human A549 cells, analyses of mouse L1475 cells upon cisplatin treatment showed increased expression of senescence mediators such as *Cdkn1a* (p21^WAF1/Cip1^), with reduced expression of genes promoting cell cycle progression (such as *E2f*, *Cdc* and *Cdk* genes) and DNA replication (such as *Pol*, *Dna2* helicase and *Ssbp1* genes). These cells also presented the highest number of uniquely upregulated genes as opposed to docetaxel-treated cells (Extended Data Fig. 3d). Characterisation of the expression of known SASP factors confirmed a general increase in the transcription of inflammatory ligands and proteases in senescent murine cells, such as *Il1a*, *Il6*, *Tgfb2*, *Tgfb3* and *Mmp3* genes (Extended Data Fig. 3e).

To identify distinctive SASP factors potentially driving increased proliferation in recipient A549 cells, we used a microenvironment microarray (MEMA) platform. This recent technology allows the screening of the functional effects of multiple combinations of individual factors and cytokines in an unbiased high-throughput manner and the subsequent analysis of treated cell read-outs through advanced imaging^21, 22^. Remarkably, analysis of the proportion of high-EdU+ A549 cells upon 72 h treatment with an array of 64 human SASP-related ligands revealed that exposure to TGFβ-2 and TGFβ- 1 led to the highest A549 proliferation rate of all factors tested in the array (Fig. 3e). Representative immunofluorescence images of the wells stained for DAPI, EdU and KRT5 (as a marker of tumour epithelial cells) can be seen in Fig. 3e, right, highlighting the increased number of EdU-positive cells upon exposure to TGFβ-1 and TGFβ-2. We therefore hypothesized that increased secretion of TGF-β ligands could be mediating the tumour-promoting effects observed in cells that are exposed to cisplatin-derived SASP. Thus, we next confirmed that cisplatin-induced senescent cells present an increased expression of *TGFB1*, *TGFB2* and *TGFB3* ligands in our RNA-seq data (Fig. 3f), which was further validated by RT-qPCR analysis in both A549 and L1475(luc) cells (Extended Data Fig. 3f). Western blot examination of control and senescent A549 cell extracts revealed increased protein levels of inactive and active TGFβ-1 in cisplatin-induced senescent cells versus control A549 cells, while free TGFβ-2 and TGFβ-3 expression was decreased (Fig. 3g). Biologically active TGFβ ligands are tightly regulated in producing cells, where they are normally stored in a latent form, while the efficient secretion, folding and extracellular deposition requires the complex regulation of multiple steps^23^. To determine whether the lower intracellular levels of the ligands observed by Western blot were due to an increased secretion compared to control cells, we analysed the levels of the active form in the SASP by ELISA. Importantly, this confirmed that the CM of cisplatin-induced senescent cells is significantly enriched in active TGFβ-1, TGFβ-2 and TGFβ-3 compared to control and docetaxel- and palbociclib-induced senescent cells (Fig. 3h).

Together, these data indicate that chemotherapeutic treatment results in markedly distinct senescent transcriptional phenotypes in human and murine lung adenocarcinoma cells, and suggest the enriched secretion of TGFβ ligands in cisplatin-induced senescent CM as potential SASP candidates promoting increased proliferation rates of recipient lung cancer cells.

### TGFBR1-driven activation of Akt/mTOR pathway orchestrates the induction of malignant NSCLC traits upon exposure to cisplatin-derived SASP

Bioactive TGFβ cytokines secreted to the extracellular space activate downstream signalling responses in recipient cells by driving the dimerization of TGFβR1 and TGFβR2, receptor serine and threonine kinases, respectively^15, 24^. To test whether TGFβR1 was involved in the observed tumour-promoting effects derived from the exposure to cisplatin-induced SASP, A549 and L1475(luc) cells were grown in the presence of control- and cisplatin-derived CMs and treated with galunisertib, a TGFBR1 inhibitor. Cell confluency and relative cell count analysis at 48 h revealed that TGFBR1 inhibition significantly hindered the increased proliferation of cells when exposed to cisplatin-derived SASPs (Fig. 4a). We next observed that galunisertib treatment also resulted in a significant reduction in the number of colonies formed after exposure to cisplatin-induced senescent A549 and L1475(luc) CM for 10 days, compared to cisplatin-derived SASP exposure alone (Fig. 4b). In addition, TGFBR1 inhibition prevented the increase in the number of spheres and sphere size observed in 3D co-cultures of A549 and L1475(luc) cells with cisplatin-induced senescent cells (Fig. 4c). To determine if the effects observed with galunisertib treatment were specific to the inhibition of TGFβR1, we silenced the expression of *TGFβR1* and *Tgfbr1* through the generation of A549 and L1475(luc) cell lines constitutively knocked down for these genes, respectively (Extended Data Fig. 4a). In agreement with the effects observed with galunisertib, reduced expression of *TGFβR1* and *Tgfbr1* resulted in a decreased impact on enhanced proliferation when A549 and L1475(luc) cells were exposed to cisplatin-derived CM, compared to parental and scrambled-transduced cells (Extended Data Fig. 4b). In addition, we also observed that treatment of A549 cells with recombinant human TGFβ1 ligand significantly increased cell growth (Extended Data Fig. 4c), further validating our observations obtained with the high-throughput MEMA platform. These results therefore confirm that the activation of TGFβR1 in recipient cells, likely through TGFβ ligands secreted by cisplatin-induced senescent cells, is responsible for the increased tumour growth driven by this particular SASP.

**Figure 4.**
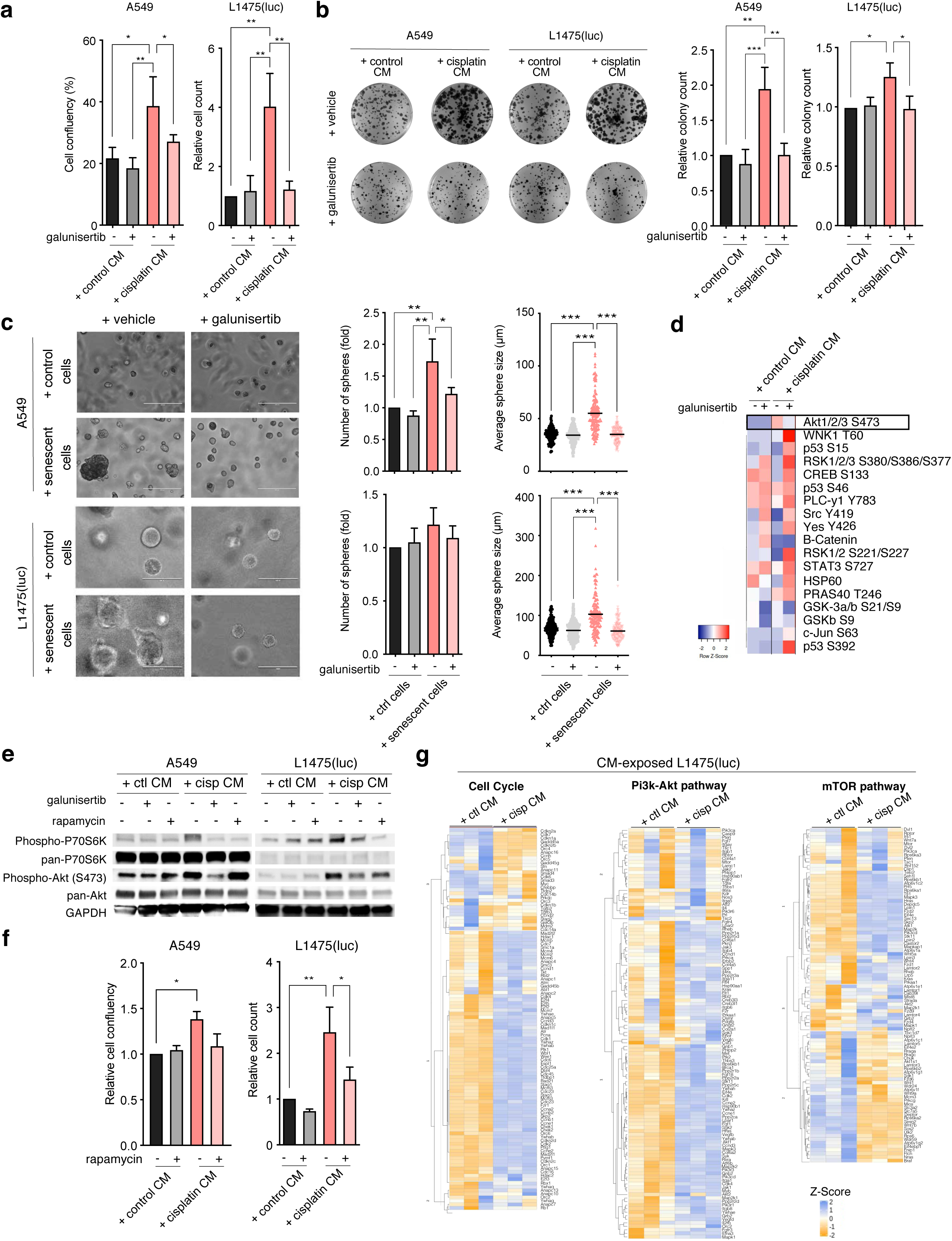
TGFβR1-driven activation of Akt/mTOR pathway orchestrates the induction of malignant traits upon exposure to cisplatin-derived SASP. **a.** Cell confluency and relative cell count of A549 and L1475(luc) cells exposed to control and cisplatin-induced senescent CM with and without 50 µM galunisertib treatment for 48 h. **b.** Left, representative images of A549 and L1475(luc) colonies formed upon exposure to control and cisplatin-induced senescent CM for 10 days with and without 50 µM galunisertib treatment. Right, quantification of the number of colonies formed relative to control condition treated with vehicle. **c.** Left, representative images of A549 and L1475(luc) tumour spheres formed after 7 days of two-phase co-culture with control or chemotherapy-induced senescent cells with and without 50 µM galunisertib treatment. Middle, number of spheres formed in each condition, relative to co-culture with control cells treated with vehicle. Right, average sphere size in each condition. For average sphere size, a total of 150 spheres from 3 independent biological repeats were measured. **d.** Heatmap of pixel intensity z-score of each kinase phosphorylation listed upon 30 min exposure to control and cisplatin-induced senescent CM in A549 cells **e.** Western blot analysis of whole extracts of A549 and L1475(luc) cells exposed to control and cisplatin-induced senescent CM for 30 min with and without 50 µM galunisertib and 1 µM rapamycin. **f.** Cell confluency and cell count of A549 and L1475(luc) cells exposed to control and cisplatin-induced senescent CM for 48 h upon 1 µM rapamycin treatment, relative to cells exposed to control CM with vehicle. **g.** Heatmap displaying expression z-scores of gene expression changes in Cell Cycle, Pi3k and mTOR pathways and their hierarchical clustering in L1475(luc) cells exposed for 12 h to control and cisplatin-induced senescent cells CM. All data represents mean ± SD (n=3), except for far-right volcano plots on panel **c**, where variation is not represented. Heatmaps represent average Z-score of 3 independent biological repeats. Statistical significance in **a**, **b**, **c** and **f** was assessed by two-tailed one-way ANOVA or two-way ANOVA followed by Tukey’s multiple comparisons test, *p < 0.05, **p < 0.01, ***p < 0.005.

Intriguingly, TGFβ cytokines can signal through the activation of several different pathways in recipient cells, which, paradoxically, can result in both tumour-suppressive and tumour-promoting effects^15, 25^. In order to uncover the mechanism involved in the transduction of TGFβR1 activation upon exposure to cisplatin-induced senescent CM, we explored protein phosphorylation changes in A549 recipient cells exposed to control and cisplatin-induced senescent CM with and without galunisertib treatment. Phospho-kinase array analysis revealed an increased phosphorylation of Akt1/2/3 at residue S473, one of the activating sites of this kinase, upon exposure of cells to cisplatin-induced senescent CM, while TGFβR1 inhibition through galunisertib treatment reduced levels of such phosphorylation (Fig. 4d and Extended Data Fig. 4d). This suggested that binding to TGFβR1 upon exposure to cisplatin-derived SASP mediates the downstream activation of Akt. As the Akt/mTOR/p70S6K is a cascade known to promote cell proliferation, we further investigated this signalling pathway. As observed in our phospho-kinase arrays, we detected an increase in phospho-Akt S473 in both A549 and L1475(luc) cells exposed to cisplatin-induced CM by Western blotting, which was diminished upon galunisertib treatment (Fig. 4e). Importantly, we observed the same trend in the phosphorylation of p70S6K, suggesting that phosphorylation of Akt results in the activation of p70S6K, a kinase downstream of mTOR that induces protein synthesis and cell cycle progression (Fig. 4e). To determine the implication of mTOR and this kinase, we treated CM recipient cells with rapamycin, a well-known mTOR inhibitor^26^. While increased phosphorylation of Akt at S473 remained unchanged in cells exposed to cisplatin-derived SASP and rapamycin, phosphorylation of p70S6K was reduced, confirming the connection of mTOR in the cascade stimulated upon TGFβR1 and Tgfbr1 activation in A549 and L1475(luc) cells, respectively, exposed to CM from cisplatin-induced senescent cells (Fig. 4e). To further validate the effect of this pathway on the phenotypic response observed upon exposure to the SASP, we exposed A549 and L1475(luc) cells to control and cisplatin-induced senescent CM and observed that rapamycin also hampered the effect on proliferation driven by cisplatin-derived SASP (Fig. 4f). Representative images of A549 cell confluency under each CM condition and upon galunisertib and rapamycin treatments are shown in Extended Data Fig. 4e.

We next performed RNAseq analyses of L1475(luc) cells exposed to control and cisplatin-derived SASP for 12 h. This confirmed the upregulation of genes involved in the cell cycle in cisplatin CM-exposed cells (such as *E2f*, *Cdc* and *Cdk* genes) (Fig. 4g), consistent with the effects observed in our *in vitro* and *in vivo* analyses. In addition, we explored changes at the transcriptional level in Pi3k- Akt and mTOR pathways, and observed a marked increase in the expression of genes involved in both pathways in cells exposed to cisplatin-derived SASP compared to control (Fig. 4g), validating the activation of these pathways upon exposure to this particular SASP in murine lung adenocarcinoma cells.

Together, these results demonstrate that the exposure of lung cancer A549 and L1475(luc) cells to the SASP derived from cisplatin-induced senescent cells results in the TGFBR1-mediated activation of the Akt/mTOR/P70S6K pathway, leading to increased cell proliferation.

### TGFβR1 pharmacologic inhibition effectively blocks pro-tumorigenic effects derived from exposure to cisplatin-induced senescence *in vivo*

To validate our mechanistic insights in *in vivo* settings, we subcutaneously transplanted cisplatin-induced senescent A549 cells together with untreated A549 cells, and subjected mice to galunisertib and senolytic treatment with ABT-737 during tumour development (Fig. 5a). As expected, the ablation of senescent cells in the tumours through senolytic treatment prevented the increase in tumour growth in co-transplanted xenografts, compared to co-transplanted tumours treated with vehicle only (Fig. 5b,c). Of note, galunisertib treatment in mice also significantly blocked the tumour-promoting effect derived from co-transplantation with senescent cells in the xenografts (Fig. 5b,c), confirming the efficiency of inhibiting TGFβR1 as a means to prevent the deleterious effects derived from cisplatin-induced senescent SASP *in vivo*. As an internal control, no impact in the progression of control A549 tumour xenografts was observed either with ABT-737 nor galunisertib treatment, suggesting that the effect is driven by transplanted cisplatin-induced senescent A549 cells.

**Figure 5.**
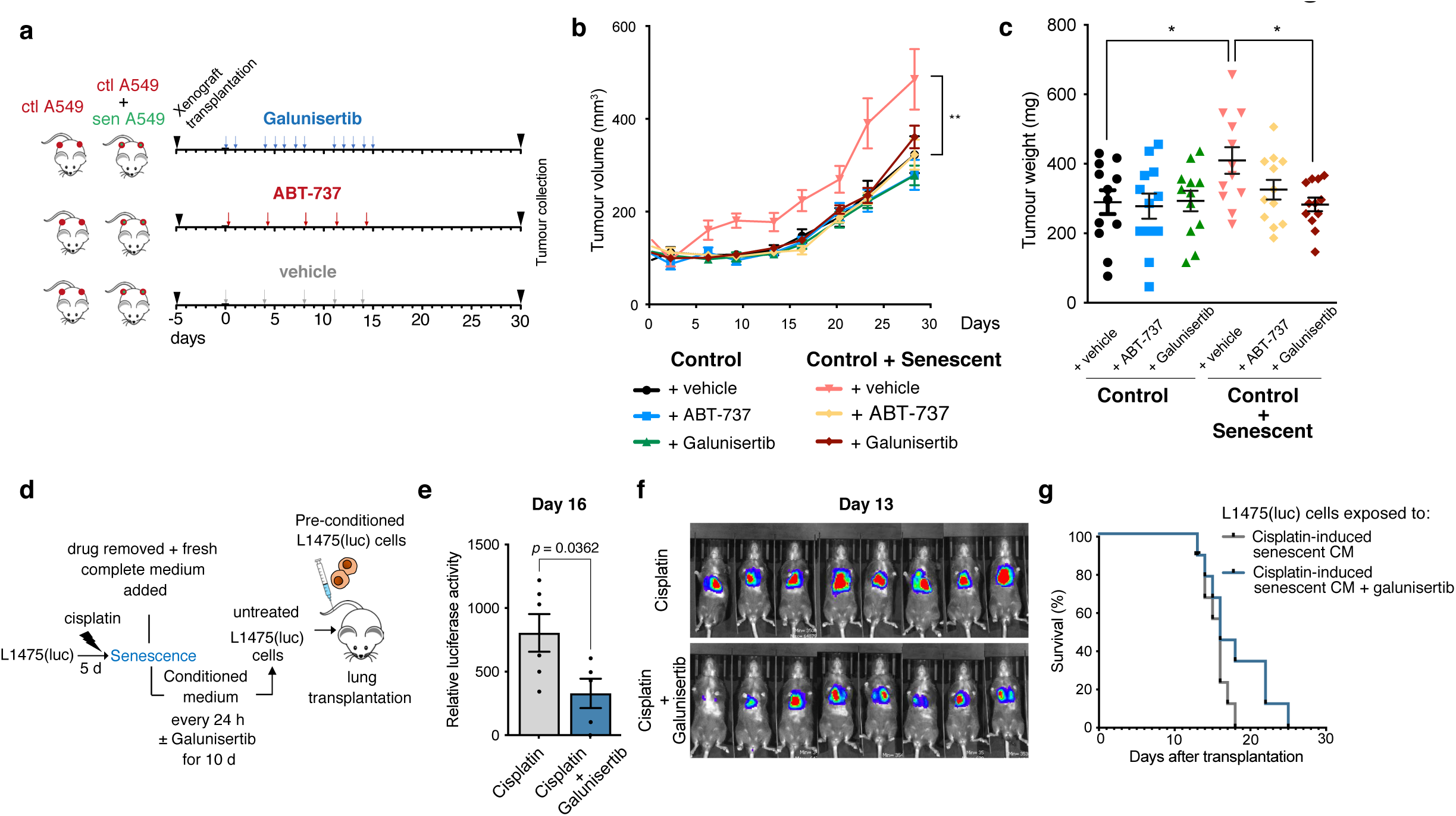
TGFβR1 inhibition effectively prevents enhanced proliferation derived from the exposure of cisplatin-induced senescent SASP in xenograft and orthotopic models of lung adenocarcinoma. **a.** Schematic representation of experimental layout. Briefly, animals were transplanted subcutaneously with either 4x10^6^ untreated A549 cells or 4x10^6^ untreated A549 combined with 1x10^6^ cisplatin-induced senescent A549 cells. At 5 days post-transplantation, animals were subjected to 150 mg/kg body weight galunisertib treatment, 25 mg/kg body weight ABT-737 or vehicle at timings depicted. Tumour volume was measured twice a week and tumours were resected at day 30 post-initiation of treatment. **b.** Tumour volume over time for each experimental condition (n=12 tumours per group). **c.** Tumour weight of each experimental group upon resection of tumours at day 30. **d**. Schematic representation of experimental layout. Briefly, L1475(luc) cells were exposed to cisplatin- induced senescent CM continuously for 10 days and treated with either vehicle or 50 µM galunisertib. Pre-conditioned cells were subsequently transplanted in the lungs of mice via tail-vein injection. Tumour burden was then assessed twice a week by bioluminescence imaging and survival was determined as the time from tumour transplantation until the onset of moderate signs of disease. **e.** Relative luciferase activity at day 16 of mice transplanted with cells pre-conditioned to cisplatin-derived CM with and without galuniseritb (n= 6). **f.** Representative images of luciferase activity at day 13 post- transplantation from each experimental group. **g.** Survival curve of mice from each experimental group. Statistical significance was determined by one- or two-way ANOVA followed by Tukey’s multiple comparisons test, and by two-tailed Student’s t-test. Survival analysis was performed using the Kaplan- Meier method and a two-sided log-rank test was conducted to determine statistical significance. *p < 0.05, **p < 0.01, ***p < 0.005, ****p < 0.0001.

We next investigated the effects of TGFβR1 inhibition simultaneous to the exposure to cisplatin-induced senescence CM using a KRas^G12D/WT^;p53^-/-^ orthotopic murine model. To this end, L1475(luc) cells were exposed to cisplatin-induced senescent CM with or without galunisertib for 10 days, and were subsequently transplanted in the lungs of C57BL/6 mice via tail-vein injection (Fig. 5d). Note that, at the tested concentrations galunisertib alone had no, or a negligible effect, on the proliferation of both L1475(luc) or A549 cells ((Fig. 4 a,b,c). Analysis of relative luciferase signal in mice at day 16 post-transplantation revealed a significantly decreased tumour burden in those animals that had been transplanted with cells simultaneously exposed to cisplatin-derived SASP and galunisertib (Fig. 5e). Representative images of luciferase activity are shown in Fig. 5f. Finally, we observed an increased survival in mice transplanted with tumours that had been simultaneously pre-treated with the Tgbfr1 inhibitor during the exposure to cisplatin-induced senescent CM (Fig. 5g).

Altogether, the above results provide *in vivo* evidence of the implication of TGFβR1 signalling in exacerbating tumour progression through cisplatin-induced senescent paracrine effects in human and murine lung cancer models.

### Platinum-based neoadjuvant chemotherapy induces tumour senescence in NSCLC patients, and results in the increased activation of the Akt/mTOR pathway

Anticancer therapeutics, including cytotoxic and genotoxic agents such as cisplatin, have been widely reported to induce senescence in experimental conditions^10, 27^. However, evidence of therapy-induced senescence in NSCLC patients, other than imperfect studies solely based on the detection of SA-β-gal in tumour specimens^2^, remains very limited. In order to further explore the potential of a senescent response upon exposure to platinum-based chemotherapy in human clinical settings, we used human samples from patients that were subjected to preoperative cisplatin-based chemotherapy followed by surgery (Fig. 6a). We analysed the expression of different proliferation and senescence-related markers in stage II-III human lung adenocarcinoma samples resected from individuals within one month of chemotherapeutic regimen completion as well as Stage II-III treatment-naïve lung adenocarcinomas (see Supplementary Table 1 for detailed clinical and pathological information). Histological analysis showed that Ki-67 detection is heterogeneous and scattered, but predominantly expressed throughout the tumours of treatment-naïve patients, compared to chemotherapy-treated specimens (Fig. 6b,c; Extended Data Fig. 5a). When compared to treatment-naïve tumour samples, cisplatin-treated tumours presented a significantly higher proportion of p21-positive cells, generally accumulating in the outer regions of the lesions and presenting a more dispersed pattern in the inner areas (Fig. 6b,c). Importantly, consecutive sections where positive signal was observed for the proliferative marker revealed a general lack of overlap with the expression of the senescent marker p21, potentially indicating the implementation of a diverse senescent response in the tumours (Fig. 6b; Extended Data Fig. 5a). Moreover, p21 areas overlapped with SenTraGor^TM^-positive (GL13) staining, which accounts for lysosomal liposfucin-positive areas and is considered an equivalent senescent biomarker of SA-β-gal activity suitable for formalin-fixed material (Fig. 6b,c)^28, 29^. The regional association between GL13 and p21 is therefore compatible with the accumulation of a cisplatin-induced senescent response across different tumour areas (see also Extended Data Fig. 6).

**Figure 6.**
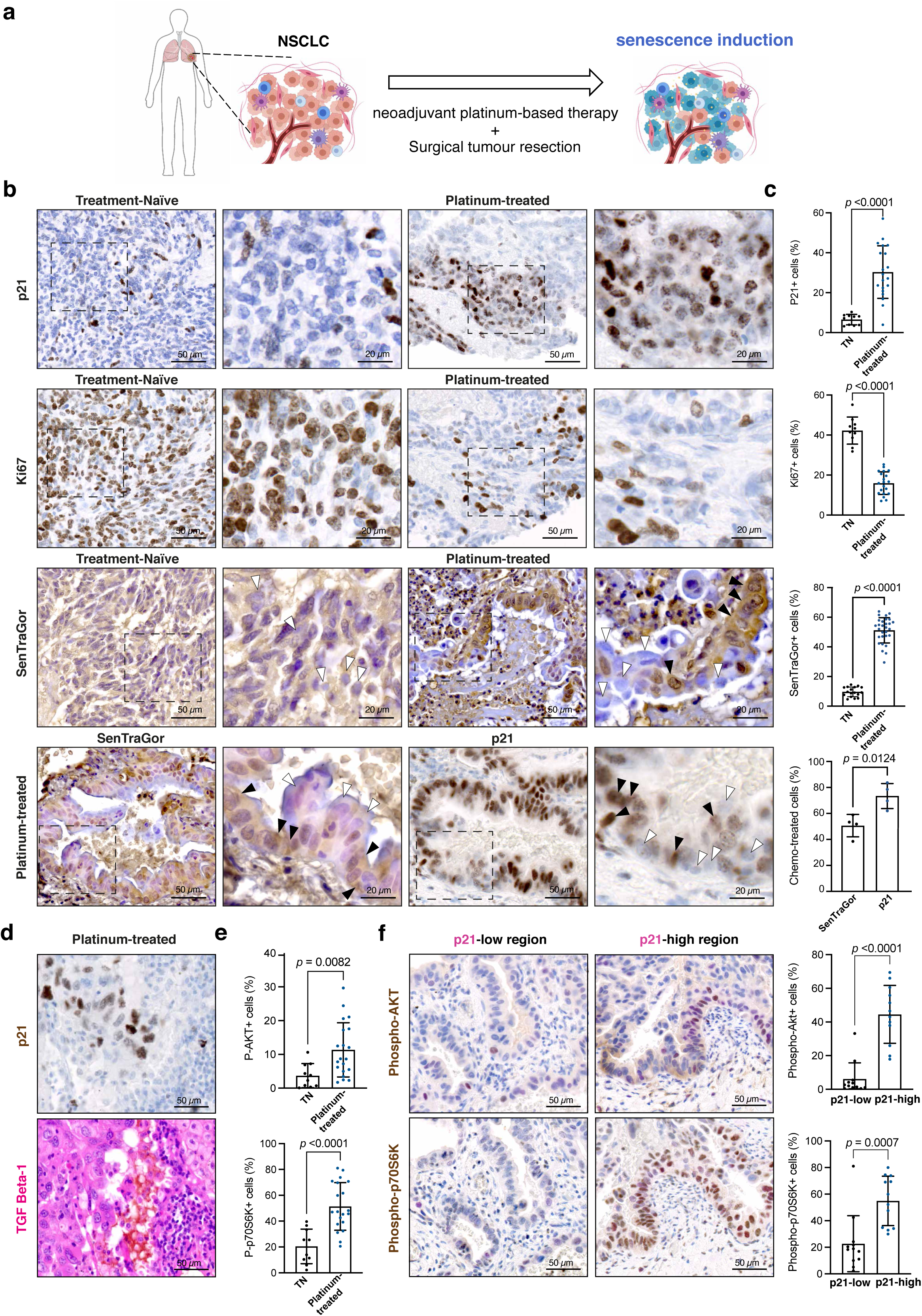
Neoadjuvant platinum-based therapy induces senescence in NSCLC patients and this correlates with increased phospho-AKT and phospho-P70S6K signalling in surrounding cells. **a.** Schematic representation of human NSCLC samples analysed in this study. **b.** Representative histological images of NSCLC specimens resected from treatment-naïve or platinum-treated patients and subjected to Ki67, p21, and SenTraGor (GL13) staining. Scale bar = 50 µm or 20 µm as depicted. Black arrowheads show SenTraGor+ or p21+ cells, white arrowheads show SenTragor- or p21- cells. **c.** Quantification of Ki67+, p21+, and SenTraGor (GL13)+ cells per total cells in treatment-naïve (TN) or platinum-treated specimens. Bottom, percentage of p21+, and SenTraGor (GL13)+ cells in consecutive sections of platinum-based treated patients. For quantification, 5 representative images were quantified per sample. **d.** Representative histological images of NSCLC specimens resected from platinum-treated patients and subjected to staining for p21 and TGFβ-1 in consecutive tissue sections. **e.** Representative histological images of NSCLC specimens resected from platinum-treated patients and subjected to co-staining for p21 (pink) and phospho-AKT or phospho-P70S6K (brown). Scale bar = 50 µm as depicted in the image. **f.** Left, quantification of phospho-AKT+ and phospho-P70S6K+ cells per total cells in p21-low and p21-high areas. For quantification, 3 representative images for each p21- low/high areas from each patient were analysed. P21-low areas were defined as those with <10% p21+ cells/total cells; p21-high areas were defined as those with >10% p21+ cells/total cells in the area. Right, quantification of phospho-AKT+ and phospho-P70S6K+ cells per total cells in treatment-naïve (TN) or platinum-treated specimens. For quantification, 5 representative images were quantified per sample. Statistical significance was determined by two-tailed Student’s t-test.

Given the evidence generated in this study linking the implementation of cisplatin-induced senescence and the paracrine activation of the Akt/mTOR pathway in a TGFβR1-dependent manner through the SASP, we also explored the expression of TGFβ ligands, phospho-AKT and phospho- P70S6K in the samples. Representative consecutive sections show the accumulation of TGFβ-1 in areas of p21-positive cells (Fig. 6d). Remarkably, these analyses showed a significantly increased reactivity against AKT and P70S6K phosphorylation within the cisplatin-treated tumours, which intriguingly correlated with the regions of accumulated p21-positive expression within the lesions (Extended Data Fig. 5a). Quantification of cells positive for phospho-AKT and phospho-P70S6K revealed that the increased phosphorylation of such kinases was significant, compared to treatment-naïve specimens (Fig. 6e), indicating an association between senescence induction and the activation of this pathway. To further determine the expression patterns and the potential link between the expression of the senescent marker p21 and the activation of these kinases, we performed co-stainings and analysed the levels of each phosphorylation in p21-low and p21-high expression regions in chemotherapy-treated tumours. Remarkably, these analyses revealed a significantly higher expression of phospho-AKT and phospho-P70S6K in regions with a marked accumulation of p21-positive cells (Fig. 6f; Extended Data Fig. 5b), indicating an association between the induction of senescence in the tumours and the proximal activation of the Akt/mTOR pathway in nearby human lung adenocarcinoma cells following platinum-based chemotherapy.

Taken together, these analyses provide supportive evidence of the induction of cellular senescence in human NSCLC tumours following cisplatin treatment, and show that accumulated senescent cells are tightly linked to the activation of the Akt/mTOR signalling in the tumour microenvironment, providing validation of the mechanistic insights of the paracrine effects of cisplatin-derived SASP in clinical settings.

### Cisplatin and galunisertib concomitant treatment reduces tumour burden and significantly enhances survival in murine models of lung cancer

Given the evidence in this study demonstrating the detrimental paracrine impact of cisplatin-induced senescent lung cancer cells on tumour progression, we next aimed to determine whether combination treatments that can prevent the deleterious non-autonomous effects of senescence accumulation in the tumours could serve as a novel and more efficient strategy to improve lung cancer treatment. To this end, we first used the KRas^G12D/WT;p53-/-^ orthotopic murine model and subjected lung tumour-bearing mice to either single treatment with cisplatin or galunisertib or a regimen combining the two pharmacologic drugs, and analysed tumour burden over time (Fig. 7a). As we have previously reported using this model, cisplatin treatment of L1475(luc)-transplanted mice results in increased SA-β-gal activity and expression of biomarkers of senescence (e.g. p21) in tumour-bearing lungs^20^. In addition, RT-qPCR analysis of RNA extracted from whole lungs upon 7 days of treatment (Extended Data Fig. 7a) revealed increased gene expression levels of the senescence marker *Cdkn1a* as well as *Tgfb1* and *Tgfb2* in cisplatin-treated animals compared to the vehicle group (Extended Data Fig. 7b).

**Figure 7.**
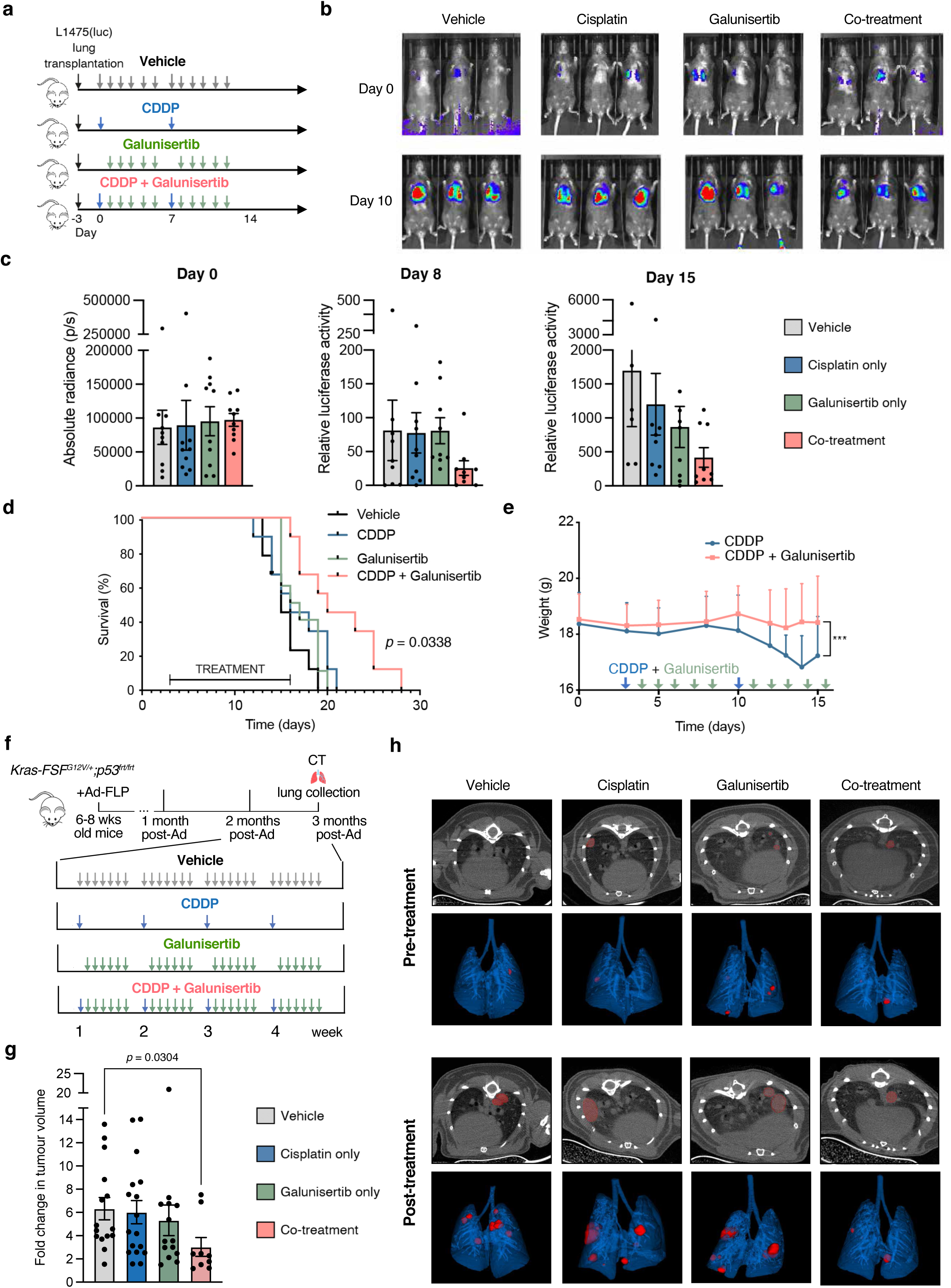
Cisplatin and galunisertib concomitant treatment reduces tumour burden and significantly enhances survival in orthotopic and KRas-driven models of lung cancer. **a.** Schematic representation of experimental layout. Animals were transplanted with L1475(luc) in the lungs via tail-vein injection, and after 3 days, they were subjected to 1.5 mg/kg body CDDP, 150 mg/kg body weight galunisertib, a combination of both or vehicle at the frequency shown in the timeline. Tumour burden was assessed twice a week by bioluminescence imaging, and survival was determined as the time from tumour transplantation until the onset of advanced signs of disease. **b**. Representative images of luciferase activity in mice at day 2 and 10 post-initiation of treatment. **c.** Absolute radiance and relative luciferase activity recorded in each experimental group at day 0, 8 and 15 post-initiation of treatment (n=9 per group). **d.** Survival curve of mice in each experimental condition. **e.** Weight of mice subjected to CDDP and CDDP + galunisertib treatment over time. **f.** Schematic representation of experimental layout. Briefly, lung tumour was induced in 6-8 week-old Kras-FSF^G12V/+^;p53^frt/frt^ animals. After 4 weeks, lungs were imaged by CT scan and number of tumours/tumour volume was analysed. Animals were subjected to 1.5 mg/kg body CDDP, 150 mg/kg body weight galunisertib, a combination of both or vehicle at the frequency shown in the timeline. 4 weeks upon treatment completion, animals were subjected to CT scanning for lung tumour burden assessment (Vehicle and CDDP groups, n=3 mice; Galunisertib and CDDP+Galunisertib groups, n=4). **g.** Fold change in tumour volume at week 12 post-Ad (4 weeks after completion of treatment) relative to initial volume measured by CT scan prior to treatment (vehicle, n=15 tumours; CDDP only, n=17; Galunisertib only, n=14; CDDP+Galunisertib, n=11). **h.** 3D rendering reconstructions of scanned lungs from each experimental group at week 4 (pre-treatment) and week 12 (post-treatment) post-Ad. Tumours are shown in red colour. Data are shown as mean ± SD. Statistical significance was assessed by ANOVA test using Welch’s correction for unequal variances in luciferase activity measurements, and by Kruskal-Wallis test in fold change tumour volume analyses. Survival analysis was performed using the Kaplan-Meier method and a two-sided log-rank test. Statistical significance in weight differences were assessed by two-sided one-way ANOVA, followed by Tukey’s comparisons test, ***p < 0.005.

Analysis of luciferase activity revealed that while single-drug treatments did not decrease luciferase activity signal compared to vehicle treatment, animals treated with the combined regimen presented a trend for decreased tumour burden as compared to the other experimental groups (Fig. 7b,c). Importantly, cisplatin and galunisertib co-treatment significantly improved median survival by 33% (20 days compared to 15 days in vehicle-treated mice; Fig. 7d). Finally, as an indirect measure of chemotherapy systemic side-effects, we studied the impact of the addition of galunisertib to cisplatin treatment on body weight throughout the therapeutic regimen, and observed that animals receiving the combination of the two drugs displayed improved tolerability and decreased weight loss and fluctuation, compared to cisplatin-only-treated animals (Fig. 7e).

Finally, we aimed to validate our observations using a distinct and more physiologically relevant model of K-RAS-driven lung adenocarcinoma, driven by endogenous expression of oncogenic Kras-G12V^30^. Mice bearing lung adenocarcinomas at 9 months post-cancer initiation were subjected to four cycles of cisplatin for one month and subsequently histological analyses of the tumours were performed (Extended Data Fig. 7c). Representative images of the tumours immunoprobed for Ki67, p21 and the DNA damage marker γH2AX, along with SA-β-gal co-staining, are shown in Extended Data Fig. 7d. This confirmed that chemotherapy significantly reduced the expression levels of the proliferative marker Ki67, while it significantly increased p21 and γH2AX, consistent with the induction of cellular senescence in the tumours (Extended Data Fig. 7e). Next, we subjected tumour-bearing Kras^+/FSFG12V^;p53^frt/frt^ mice ^30, 31^ to 4 cycles of cisplatin-based chemotherapy, alone or in combination with galunisertib treatment, as shown in Fig. 7f. Tumour burden analysis upon CT imaging of murine lungs demonstrated that the combined administration of cisplatin with the TGFβR1 inhibitor resulted in a significantly decreased fold change in tumour volume, compared to animals treated with vehicle or single treatments (Fig. 7g). Representative CT scans and 3D reconstructions of tumour-bearing lungs from mice in each experimental group before and after treatment are shown in Fig. 7h.

Taken together, our results emphasize the importance of modulating tumour-promoting activities of therapy-induced senescence and reveal the clinical potential of combinatory treatments based on platinum chemotherapy and TGFβR1 inhibitors to improve patient outcomes in lung cancer management.

## Discussion

Platinum-containing chemotherapy regimens remain standard treatments for many NSCLC patients and deliver significant patient benefit, either alone or in combination with immunotherapy^1^. Despite advances over the past decade, including therapies targeting oncogenic kinase addiction (present in a small subset of patients), anti-PD(L)1 immunotherapy and chemo-immunotherapy combinations, the long-term outcomes for most patients with advanced disease remain poor, with treatment resistance and progression being essentially inevitable^32, 33^. Prolonged responses to these treatments are disappointingly infrequent, particularly in advanced disease^1^. Treatment of early-stage NSCLC by surgical resection and adjuvant platinum-containing chemotherapy confers a better prognosis but a significant percentage of these patients relapse due to unrecognised micrometastatic disease at presentation. Therefore, further insights into the tumour therapy response are required to develop more effective drugs and combination therapies to expand the clinical benefit to a broader patient population, and hence to improve clinical outcomes in NSCLC. Early studies using human NSCLC specimens after neoadjuvant platinum-based therapy reported the induction of SA-β-gal activity affecting both tumour areas and the background lung^2^, although further assessment of senescence and mechanistic insights remained elusive. Here, we show that a cisplatin-induced SASP enriched for TGF-β ligands strongly promotes cancer cell malignant traits and tumour progression in a variety of lung cancer mouse models, a process driven by the activation of TGFβR1 signalling and the Akt/mTOR pathway in recipient lung cancer cells. Of translational relevance, we provide mechanistic validation *in vivo* and in NSCLC patients subjected to preoperative neoadjuvant platinum-based therapy, and also show that concomitant treatment with cisplatin and galunisertib, a TGFβR1 inhibitor, significantly reduces tumour burden and increases survival in murine models of lung cancer.

Senescence is a well-known tumour suppression mechanism arresting premalignant and proliferative cancer cells but increasing evidence shows that the persistence of chemotherapy-induced senescent cells and their SASP phenotypes can also yield opposing outcomes driving tumour relapse, treatment resistance, metastasis and a variety of chemotherapy-related adverse effects^34^, although the impact of treatment-induced senescence in the particular case of NSCLC remains poorly understood. These unwanted and potentially detrimental effects can apply to both senescent cancer cells and senescent stromal cells, such as fibroblasts^8, 18, 35–38^. We examined here the effect of the SASP from senescent lung cancer cells induced by different types of chemotherapies in both human and murine lung models, and we demonstrate that the SASP derived from cisplatin treatment in lung adenocarcinoma cells drives enhanced proliferation, migration, EMT and maximal respiratory abilities of recipient lung cancer cells, in contrast to SASPs from cells exposed to docetaxel and palbociclib. Previous reports have reported similar effects derived from doxorubicin-induced senescence in breast cancer and melanoma cells^18, 39, 40^, and have even explored the effect of cisplatin on A549 lung adenocarcinoma cells, but the treatment conditions used by the authors were suboptimal for the effective induction of cellular senescence^39^. In addition, we found that increased proliferation rates of lung cancer cells also occur upon cell exposure to a carboplatin-induced SASP and this finding was expanded to a human ovarian cancer cell line. Platinum-based drugs (similarly to doxorubicin) are genotoxic agents that induce a robust DNA damage response, while docetaxel blocks microtubular depolymerisation and palbociclib directly inhibits Cdk4/6. It has been postulated that persistent DNA damage prompts an enhanced and proinflammatory SASP^37^, and therefore it is conceivable that the divergent responses to the different SASP cocktails induced by these three chemotherapeutics are explained by the interplay between the signalling cascades activated upon cell exposure to the specific drug and the different regulatory pathways of the SASP.

Importantly, in this study we show that cisplatin-induced senescence increased tumour growth in A549 tumour xenograft and L1475 KP orthotopic models of lung cancer through paracrine effects on untreated cancer cells. In further support of tumour-promoting activities of platinum-induced senescence, we show that senolytic ABT-737 treatment reduces tumour progression only in mice co-transplanted subcutaneously with cisplatin-induced senescent and untreated A549 cells but not in A549 only xenografts, and that the concomitant treatment of Kras^FSFG12V/+^ lung tumour-bearing mice with cisplatin and ABT-737 effectively decreases tumour burden and blocks progression compared to single agent cisplatin treatment. Remarkably, these observations are in accordance with early findings reporting that NSCLC patients subjected to platinum-based therapy and showing in situ tumour activity of SA-β-gal have a significantly reduced survival when compared to their SA-β-gal negative activity counterparts^41^, which highlights/emphasizes the potential detrimental impact of unresolved (i.e. uncleared) senescence driven by platinum-based treatment and suggests its assessment as a predictor of lung cancer treatment outcomes and/or patient stratification. Our experiments also provide preliminary evidence that platinum-based therapy results in enrichment of senescence biomarkers in the functional lung pre-tumour induction, and some detrimental effects, including increased tumour burden and reduced survival, are exacerbated in adult/aged mice when compared to young mice. This observation correlates with higher levels of potential toxicities of chemotherapies in aged individuals^42^.

We establish that the SASP secretome derived from cisplatin treatment of lung adenocarcinoma cells is enriched in TGF-β ligands, which was validated by transcriptomic analyses (including RT-qPCR and RNAseq gene expression profiles), and proteomic analyses (including MEMA, protein arrays and ELISA assays with CM) by using human A549 and murine L1475 KP lung cancer cells. TGF-β is a highly pleiotropic cytokine that has multiple and dual effects in cancer, highly dependent on the extracellular context and the phenotype of the recipient cell^15^. While TGF-β often functions as a tumour suppressor in pre-malignant cells it can also act as tumour promoter in overtly malignant cells^25^. Given the multifaceted nature of this ligand and the complexity of the senescent phenotype, the relationship between the two remain largely elusive. Recent findings suggest that senescent-derived secretion of TGF-β exacerbates age-related disorders including Alzheimer’s disease, muscular atrophy and obesity^43^. However, the most prominent reported function of TGF-β at the interplay of senescence and cancer is its cytostatic effect. In fact, TGF-β has been shown to induce cellular senescence and reinforce the proliferative arrest in an autocrine fashion mainly through the Smad-driven activation of CDK inhibitors^14, 44, 45^. On the contrary, the exact molecular mechanisms underlying TGF-β-mediated promotion of cell proliferation and survival remain unclear, but it is believed that they rely on the balance of all signalling inputs received by the cell, the insensitivity to the TGF-β-induced antiproliferative cascade and the abundance or activity of TGF-β in the extracellular matrix and the tumour microenvironment.

To our knowledge, we report for the first time the mechanistic connection between chemotherapy-induced senescence, TGFβR1 signalling and stimulation of the Akt/mTOR/p70S6K pathway in lung tumours, ultimately resulting in enhanced proliferation rates of NSCLC cells. This is inferred from the observed attenuation of proliferation rates and malignant traits of lung cancer cell lines exposed to cisplatin-derived SASP upon pharmacologic inhibition of TGFβR1 (with galunisertib) and mTOR (with rapamycin). Furthermore, TGFβR1 pharmacologic inhibition with galunisertib effectively hampered the tumour-promoting effects derived from exposure to cisplatin-induced senescence in A549 tumour xenografts and orthotopic transplantation models of lung cancer. This non-canonical TGF-β signalling was first reported in 2007^46^, and several studies have since demonstrated that the interplay between TGF-β and Akt/mTOR pathways is a strong promoter of EMT and metastasis in malignant cells^47^. Indeed, the increased migratory properties and formation of protuberances during sphere development of A549 cells observed in our study suggest the acquisition of an EMT phenotype in response to cisplatin-derived SASP, an intriguing finding warranting further investigation.

Importantly and in support of these mechanistic insights, we show an association between the accumulation of cellular senescence markers (SenTraGor, p16 and p21) and the activation of the Akt/mTOR pathway in human lung adenocarcinoma samples obtained from patients that had recently received platinum-based neoadjuvant therapy. This is evidenced from extensive areas showing phospho-AKT and phospho-P70S6K staining nearby cells expressing senescence biomarkers. Interestingly, a study of stage III NSCLC patients subjected to platinum-based therapy found significantly shorter tumour doubling times (i.e. accelerated regrowth of tumours) in the interval period between the end of induction platinum-based chemotherapy and the start of radiotherapy when compared to non-chemotherapy treated patients^48^. This outcome fits with the observation that NSCLC patients subjected to neoadjuvant platinum-based therapy and showing in situ tumour expression of SA- β-gal have a significantly reduced survival when compared to their SA-β-gal negative activity counterparts^41^. While further and more detailed analyses are needed, these results are consistent with increased proliferation rates and tumour promoting activities associated with platinum-based therapies after their initial anticancer properties.

After validating the signalling pathways involved in increased tumour-promoting activities in the context of cisplatin-induced senescence we focused on the therapeutic potential of targeting such pathways in preclinical models of NSCLC. Of translational relevance our data show that concomitant treatment of cisplatin and galunisertib reduces tumour burden and significantly enhances survival in orthotopic and KrasFSF^G12V^-driven lung cancer mouse models. Galunisertib is a promising oral small molecule inhibitor currently under extensive clinical investigation both as a monotherapy and in combination with standard anti-cancer regimens for the treatment of various cancer types, including hepatocellular carcinoma, glioblastoma, ovarian cancer and pancreatic cancer, among others, and has significant potential for suppressing tumour progression and promoting immune-surveillance^49^. In addition to the TGF-β-stimulating proliferative effects in NSCLC reported herein, TGF-β is a potent immune regulator, and its circulation as part of the SASP has been shown to result in immunosuppression via the recruitment of M2 macrophages following irradiation-induced senescence in murine kidneys^50^. Although effects on immunomodulation have not been explored in our study, it is tempting to speculate that TGF-β-rich cisplatin-derived SASP might also contribute to additional adverse effects through the promotion of immunosuppression in tumours.

Alongside the focus here on the paracrine effects of chemotherapy-induced senescence, it is important to note that the SASP may not be the only contributor to treatment failure following the induction of senescence. Contrary to the long-standing belief that senescence is an irreversible state, recent research provided evidence that chemotherapy-induced senescent cancer cells have the potential to escape the stable cell cycle arrest and re-enter the cell cycle again with aggressive proliferation rates, enhanced malignancy^51, 52^ and increased stemness properties^53, 54^, thereby driving cancer relapse. Notably, chemotherapeutic treatment has recently been reported to induce very heterogenous responses across lesions and within tumours^55, 56^, and thus it is likely that a combination of different cellular processes occurs in response to the treatment, including senescent, apoptotic and resistant cancer cells as well as senescence-escapers. Indeed, a high mutational burden landscape has been linked to greater heterogeneity in tumours and poorer prognosis after first-line chemotherapy^56^. In this scenario, it is conceivable that the detrimental effects of the SASP are exacerbated, and therefore a complete and thorough dissection of tumour response in patients undergoing chemotherapy may be important to establish to optimise and personalise anti-cancer regimens.

Despite significant advances, the overall cure and survival rates for NSCLC patients remain poor, and therefore novel and more efficient treatment modalities are still needed to maximize beneficial outcomes in the clinic. Taken together, our data demonstrate that cisplatin treatment in human and murine lung adenocarcinoma cells drives the acquisition of tumour-promoting phenotypes in a paracrine manner through the implementation of a SASP enriched in TGF-β ligands. This process is driven, at least in part, by the activation of TGFβR1 signalling and the Akt/mTOR pathway in recipient lung cancer cells, and these mechanistic insights were validated in mouse models and in NSCLC patients subjected to preoperative neoadjuvant platinum-based therapy. We demonstrate that the detrimental effects of cisplatin-induced senescence can be effectively prevented in preclinical models through TGFβR1 inhibition and senolytic treatment. Finally, we propose the combination of cisplatin chemotherapy and TGFβR1 inhibitors (+/- agents with distinct mechanisms of action including anti- PD-(L)1 immunotherapy) as a novel and efficient therapeutic approach for the management of NSCLC in order to prevent the detrimental effects from unresolved bystander senescence in tumours.

## Methods

### Cells and Reagents

For *in vitro* experiments, human lung adenocarcinoma A549 cell line was obtained from ATCC and maintained in DMEM growth medium supplemented with 10% Foetal Bovine Serum (FBS). The heterozygous KRas^G12D/WT^ murine L1475(luc) cell line was generated from Kras^G12D/WT;^p53^Fx/Fx^ mice^57, 58^ and maintained in F12/DMEM growth medium supplemented with 10% FBS. The ovarian cancer cell line PEO4 was kindly gifted by Prof James D Brenton (CRUK Cambridge Institute) and maintained in RMPI growth medium supplemented with 10% FBS. All cells were grown at 37°C and 5% CO2 and were routinely authenticated and tested for Mycoplasma contamination.

For the induction of cellular senescence *in vitro*, A549 were treated with 15 µM cisplatin (CDDP, Sigma-Aldrich), 7.5 µM carboplatin (Sigma-Aldrich), 100 nM docetaxel (Sigma-Aldrich) and 15 µM Palbociclib (Cambridge Bioscience) for 7 days. L1475(luc) cells were treated with 3 µM CDDP, 100 nM docetaxel and 30 µM Palbociclib for 5 days. PEO4 cells were treated with 2.5 µM CDDP and 10 µM carboplatin. For proliferation and colony-formation assays, A549 and L1475(luc) cells were treated with 50 µM Galunisertib (Stratech) and 1 µM Rapamycin (Stratech). A549 cells were also exposed to rhTGF-beta1 ligand (Bio-Techne) at a concentration of 10 ng/mL.

### Human Biopsies and Ethical Regulations

Human lung adenocarcinoma samples were obtained from the Royal Papworth Hospital Research Tissue Bank (RPHRTB) following review by the RPHRTB project review committee (Project Number T02722). RPHRTB has derogation under the UK Human Tissue Authority (HTA) to supply samples (HTA number 12212), surplus to clinical need, that have been collected using Research Ethics Committee approved RPHRTB consent. Patients signed the RPHRTB general consent and accepted the use of their biopsy for research purposes. Information about the human samples used can be found in Supplementary Table 1.

### Animal work and in vivo treatment studies

For senescence induction upon chemotherapeutic treatment and evaluation of chemotherapy and galunisertib treatments *in vivo*, lung tumours were generated in Kras-FSF^G12V/+^ ^30^ and Kras-FSF^G12V/+;^p53^frt/frt^ ^30, 31^ mice through intranasal or intratracheal administration with FLP-expressing adenovirus. Briefly, 8- to 10-week old mice were treated once with Adeno-FLP particles (2.5x10^7^ PFU/mouse of virus) after anaesthesia (isoflurane inhalation). For virus preparation, viral particles were diluted in DMEM, precipitated with 10 mM CaCl_2_ and incubated for 20 min prior to nasal inhalation or intratracheal administration. In Kras-FSF^G12V/+^ mice, at 9 months post-tumour induction, mice were treated with 4 doses of 1.5 mg/kg body weight CDDP once weekly. In Kras- FSF^G12V/+;^p53^frt/frt^ mice, mice were subjected to either vehicle or 1.5 mg/kg body weight CDDP once weekly for 4 subsequent weeks, followed by five daily administrations of Galunisertib (LY2157299; 150 mg/kg body weight) after each CDDP dose.

For xenograft experiments, SCID mice were used. Cell suspension was prepared by mixing Matrigel (Corning) 1:1 with either 1x10^6^ control A549 cells, 1x10^6^ control and 1x10^6^ cisplatin-induced senescent A549 cells or 4x10^6^ cisplatin-induced senescent A549 cells alone. Cells were injected subcutaneously and xenograft growth was monitored using a digital calliper twice a week. 5 days post- transplantation, animals were treated with 150 mg/kg body weight Galunisertib (LY2157299), 25 mg/kg body weight ABT-737 or vehicle following the experimental timing detailed in the respective figures. Tumour volume was calculated as Volume = (D x d^2^)/2, where D and d refer to the long and short tumour diameters, respectively. Experiments were terminated at either pre-established time endpoints or when tumours reached an average diameter of 1.5 cm.

For orthotopic transplantation of lung cancer cells, C57BL/6J female mice were sublethally irradiated at 4 Gy using a Caesium source irradiator 6 h prior cell injection. Luciferase-expressing Kras^G12D/WT^ murine lung tumour L1475(luc) cells were injected intravenously (2x10^5^ cells/mouse). Baseline luminescence was recorded 24 h after transplantation, and tumour growth was monitored twice a week by bioluminescence imaging following intraperitoneal injection with D-luciferin (150 mg/kg body weight, Perkin Elmer) using an IVIS Spectrum Xenogen imaging system (Caliper Life Sciences) and Living Image® software (version 4.7.3). Relative luciferase activity was calculated as the change from baseline at indicated time points, normalised to a blank control (luciferase-negative animal). Tumour survival represents the onset of moderate signs of disease and/or 15% weight loss.

All experiments were approved for Ethical Conduct by the Home Office England and Central Biomedical Services (CBS), regulated under the Animals (Scientific Procedures) Act 1986 (ASPA), as stated in The International Guiding Principles for Biomedical Research involving Animals.

### Micro-Computed Tomography (CT) Imaging and Analysis

Mice were anaesthetized by isoflurane inhalation and scanned using a microPET-CT scanner (Mediso Medical) and Nucline™ Nanoscan software (Version 2.0) with an X-ray energy of 35 kVp, an exposure time of 450 ms, a total number of projections of 720 and one projection per step following a semi-circular single field of view scan method. Scans were reconstructed through Butterworth filter and a small voxel size. 3D reconstructions and analyses were performed using Slicer 4.10.2 software (3D Slicer). To determine tumour size, the length (longest diameter, L) and width (diameter perpendicular to the length, W) were measured and the ellipsoid formula was applied (Tumour volume = (4/3) x PI x (W/2)^2^ x (L/2)).

### In Vitro Assessment of Malignant Traits

#### Conditioned media preparation and analysis

To evaluate the effect of the different SASPs on cell behaviour in a paracrine manner, cells that had been under chemotherapeutic treatment as specified above were thoroughly washed with PBS twice and then either FBS-free- or 10% FBS-medium was added. After 48-72 h of incubation, conditioned medium (CM) from senescent cells was collected and centrifuged for 10 min at 2,500 rpm before adding onto non-senescent cells for further assessment. CMs used on all assays were freshly conditioned and were not subjected to storage/freezing.

#### Proliferation assays

For proliferation assessment, 20,000 A549 cells were seeded in a 24-well plate and let to attach overnight. The next morning, media was removed, cells were washed twice with PBS and fresh CM was added onto cells. Cells were let to grow over a period of 72 h and pictures were taken every 2-3 h with an IncuCyte® S3 Live Cell Analysis System microscope (Essen Bioscience). Cell confluency was analysed for each time-point using the IncuCyte ZOOM^TM^ software analyser (v2016B). A total of 50,000-100,000 L1475(luc) cells were seeded in 6-well plates and the next morning, cells were washed with PBS and CM from senescent plates was added onto the plates. At specified time- points, cells were trypsinised and 50 µL of Precision Count Beads™ (BioLegend) were added to each tube for cell count by flow cytometry using a BD LSRFortessa™ Cell Analyzer (BD Biosciences). Data was recorded using FACSDiva^TM^ software (v8.0.1) and analysed with FlowJo Software (v10.8.1).

#### Scratch-wound Cell Migration Assay

A total number of 50,000 cells were seeded in a 96-well plate and when cells reached 100% confluency, the WoundMaker® tool (Essen Bioscience) was applied to create a horizontal wound on each well. Cells were washed once with PBS and then CM was added onto each well. Cells were allowed to migrate over a period of 42 h and pictures were taken every 2 h with an IncuCyte® S3 Live Cell Analysis System microscope (Essen Bioscience). Relative wound confluency was analysed for each time-point using the IncuCyte ZOOM^TM^ Software analyser.

#### Colony-formation assay

A total of 500 cells/well were seeded in 6-well plates and let to attach overnight. The next morning, cells were carefully washed and CM from senescent and control cells was added to the cells. Fresh CM was replaced every 24-48 h and colonies were let to grow for 10-14 days. At the end of the experiment, colonies were washed with PBS and fixed with 4% PFA for 10 min. They were permeabilised in ice-cold methanol for 20 min, dried and stained with 0.2% crystal violet (ACROS organics) in 20% methanol for 30 min. Plates were washed with distilled water, colony plates were scanned and number of colonies were analysed using FiJi Software (v2.1.0/1.53c).

#### Low Attachment Sphere Forming assay

Cover glasses were placed inside 6-well plates and 0.01% polylysine was added and allowed to set for 30 min. A total of 2,000 cells/well were resuspended in collected CM and 10 ng/mL rhFGF (R&D Systems), 10 ng/mL hEGF (Invitrogen) and 1% N-2 Supplement (Life Technologies) were added. Cells were seeded onto coverslips and allowed to form spheres for up to 14 days. Supplemented CM was replaced every 48 h, and at the end of the experiment, spheres were fixed with 4% PFA for 20 min and imaged with an AxioScan.Z1 microscope and Zen Blue (v2.6) at 10X. Sphere number and shape was analysed using FiJi software.

#### 3D Matrigel co-culture assay

For 3D direct co-culture experiments, a total of 5,000 non-senescent cells were plated in the middle of a 24-well plate immersed in Matrigel (Corning). After matrix solidification for 20 min at 37°C, 15,000 senescent cells were carefully seeded surrounding the solidified 3D droplet. Spheres were allowed to grow for 5 to 7 days; representative pictures were taken with an EVOS Cell Imaging System microscope (ThermoFisher Scientific) and analysed using FiJi software.

### Cell Mitochondrial Stress Test

To assess oxygen consumption rate (OCR) and extracellular acidification rate (ECAT) of senescent, non-senescent and SASP-recipient cells, a total of 40,000 were seeded in an XFe24 cell culture microplate. Once attached, fresh media or CM was added (supplemented with 25 mM glucose, 1 mM pyruvate and 4 mM glutamine). Cells were incubated for 2 h at 37°C with atmospheric CO_2_ in a non-humidified incubator. Previously calibrated Seahorse cartridges were loaded with 2 µM oligomycin, 1 µM carbonyl cyanide-p-trifluoromethoxyphenylhydrazone (FCCP), 1 µM rotenone and 1 µM antimycin A. Cartridge was placed onto microplate containing cells and inserted into a Seahorse XF-24 Flux Analyzer (Agilent Technologies). Three measurements of 2 min-mix, 2 min-wait and 4 min-measure were carried out at basal condition and after each drug injection into the wells. After measurements, cells were washed with PBS and protein was extracted with RIPA lysis medium for quantification. OCR and ECAR values were normalised to total µg of protein per well.

### Gene Expression Analysis (RNA extraction and qPCR)

For gene expression analyses, RNA was extracted using the RNeasy Mini Kit (Qiagen) and where applicable, cDNA was synthesized with the High-Capacity RNA-to-cDNATM Kit (Thermo Fisher Scientific) following manufacturer’s instructions.

Gene expression was measured by quantitative real-time PCR performed on a QuantStudio Thermocycler (Applied Biosystems) using QuantStudio™ Design & Analysis Software (v1.5.1) following Luna® Universal qPCR Master Mix (New England Biolabs) protocol and amplification parameters, and using pre-designed KiCqStart® SYBR Green Primers (see details in Supplementary Table 2). Relative quantification was carried out using 2- ΔΔCt methodology.

For bulk RNA sequencing, RNA samples were assessed for optimal quality using the Agilent RNA ScreenTape System (Agilent Technologies) and submitted to BGI Sequencing Services for library preparation and sequencing (75x2 reads). Reads were aligned to the reference genome (either hg38 for human data or mm10 for mice data) using STAR (version_2.6.1d) in two-pass mode following STAR best practices and recommendations^59^. The quality of the data was evaluated using STAR (version_2.6.1d)^59^ and samtools (version 1.9)^60, 61^. PCR duplicates were removed from aligned bam files using samtools (version 1.9)^61^. Read counts were extracted from the aligned bam files using subread’s FeatureCounts (version 1.6.4)^62^. Normalisation of read counts for analysis was done according to EdgeR recommendations using the Ratio of the Variance method which accounts for inter-sample variance and the differential expression analysis of the normalised read counts between the sample groups was performed following best practices and recommendations of EdgeR^63–65^ and Limma^66^ on R environment (version 3.0.6). Pathway over-representation analyses were performed as implemented in KEGG (KEGGREST version 1.24.1)^67^, GO (GOSEQ version 1.36.0)^68^ and GSEA (FGSEA version 1.10.1)^69–71^ R packages. RNA-seq raw data are available in the EMBL’s European Bioinformatics Institute (EBI) database: European Nucleotide Archive (ENA) repository Accession Number PRJEB52271 (https://www.ebi.ac.uk/ena/browser/home) and code used for data analyses are available upon request.

### shRNA Knockdown

For TGFBR1 and Tgfbr1 knockdowns, viral particles were produced in 293T cells upon transfection with pCMVAR8.91, pMD2.G and shRNA plasmids mixed at a 1:2:3 proportion, respectively, with Lipofectamine 2000 (ThermoFisher Scientific) in Opti-MEM medium (ThermoFisher Scientific). Cells were incubated with the mix overnight and fresh medium was added the next morning. After 48 h incubation, supernatant was collected and filtered. Viral particles were added onto A549 and L1475(luc) cells, and positive selection was performed with puromycin treatment for 5 days. RNA was then extracted as described above, and lines with the highest knockdown were used for further assessment and experiments. Clone IDs, target sequence and further details of the shRNA constructs used can be found in Supplementary Table 3.

### Western Blotting

Protein was extracted using Radioimmunoprecipitation Assay (RIPA) buffer (Sigma-Aldrich) supplemented with 1 mM EDTA, cOmpleteTM EDTA-free EASYpak protease inhibitor cocktail (Roche) and PhosSTOPTM EASYpak phosphatase inhibitor cocktail (Roche). Lysates were incubated on ice for 15 min and centrifuged at 15,000 for 15 min. Protein concentration in supernatant was quantified using the PierceTM BCA Protein Assay Kit (ThermoFisher Scientific). A total of 30 µg of protein per sample were diluted in Laemmli Sample Buffer (Bio-Rad) and run for 50 min at 120V into a Mini-PROTEAN® TGX Precast Gel. Proteins were electrotransferred from the gel onto a PDVF membrane by wet tank transfer overnight at 4°C. Membrane was washed with TBS-T buffer (Tris Buffered Saline with 1% Tween 20) and blocked in 5% milk solution. Membranes were incubated with primary antibodies overnight at 4°C, washed three times with TBS-T buffer and incubated with HRP-conjugated secondary antibodies for 1 h at room temperature. Finally, membranes were incubated with Enhanced Chemiluminescence Detection Solution (ECL, Amersham) and imaged using a Xograph Compact X4 automatic processor or a ChemiDoc Imager (Bio-Rad) using Image Lab™ (v6.1). Full list of antibodies used for WB can be found in Supplementary Table 4.

### ELISA

Senescent and control cells were washed and FBS-free fresh media was conditioned for 72 h. CM was then collected, centrifuged for 10 min at 2,500 rpm and immediately stored at -80°C for ELISA analysis of TGFB ligands. Prior to the assay, ligands were activated by incubating the samples with 1 N HCl for 10 min and neutralizing them with 1.2 N NaOH/0.5 M HEPES solution. Samples were assayed immediately after using the Quantikine® Human TGF-β1 Immunosasay Kit (R&D Systems), the Human TGF-beta 2 Quantikine ELISA Kit (R&D Systems), and the Human TGF-beta 3 DuoSet ELISA Kit (R&D Systems).

### Phospho-Kinase Array

Changes in kinase phosphorylation upon exposure to the SASP was assayed using the Proteome Profiler Human Phospho-Kinase Array (R&D Systems). Briefly, A549 cells were exposed to FBS-free CM from control and senescent cells (with either vehicle or galunisertib) for 30 min at 37°C. Protein was extracted immediately after as described above and protein concentration was calculated following the Pierce^TM^ BCA Protein Assay Kit (ThermoFisher Scientific). A total of 600 µg were immediately used for the assay following manufacturer’s indications. Membranes from the kit were incubated with ECL (Amersham) and X-ray film was exposed to 1 and 10 min. Pixel intensity was calculated using FiJi software (v2.1.0/1.53c) and normalised again loading controls.

### MEMA

The MicroEnvironment MicroArray (MEMA) platform^21, 22^ was used as a high-throughput technology to determine the impact of different SASP ligands on A549 cell proliferation. Briefly, MEMA plates were blocked for 20 min with 1% non-fouling blocking agent and rinsed three times with PBS. A total of 3 x10^4^ A549 cells/well were seeded in DMEM medium containing 0.1% FBS. After 18 hours of adhesion, each well was supplemented with an experimental SASP ligand. Cells were allowed to grow for up to 72 h and were then incubated with 10 µM EdU for 1 h prior to fixation. Next, cells were fixed with 2% PFA for 15 min at RT and stored at 4°C in PBS. Cells were then permeabilised with 0.1% Triton X-100 for 15 min, washed with PBS, washed with 0.05% Tween 20 PBS (PBS-T), and incubated with Click-iT EdU detection reaction reagents (Click-iT^TM^ EdU Cell Proliferation Kit (ThermoFisher Scientific)) for 1 h while protected from light. After quenching, cells were rinsed with PST-T and subjected to immunofluorescence histochemistry (IHC) by using anti-KRT5 and anti- KRT19 antibodies (Supplementary Table 4). Cells were then washed again with PBS-T, stained with secondary antibodies, washed with PBS-T, washed with and stored in PBS, and imaged on an automated imaging system. Segment cells and intensity were calculated using CellProfiler. R-environment with custom code was used to normalize, correct variations and summarise the raw Cell Profiler derived data for each condition. Custom code and raw data is accessible on GitHub (https://github.com/markdane/A549_low_serum/tree/main/R) and Synapse (ID: syn27665118).

Full list of ligands used for MEMA experiments, uniport ID and concentrations are found in Supplementary Table 5.

### SA-β-Gal Staining

Upon senescence induction, cells were washed with PBS, and fixed and stained for Senescence Associated β-Galactosidase Activity (SA-β-Gal) using the for Senescence β-Galactosidase Staining Kit (Cell Signaling) following manufacturer’s instructions. Stained cells were imaged using an Olympus Compact Brightfield Modular Microscope (Life Technologies) and ZEN Blue (v2.6). Resected lungs from experimental mice were also stained whole-mount for SA-β-Gal following the same procedure and kit.

### SenTraGor Staining

SenTraGor staining was performed as described previously^28, 29, 72^. Whole slides were digitalised using an AxioScan Microscope Slide Scanner (Zeiss) using polarised light, and ZEN Blue Software (v2.6.) was used for image acquisition. The mean percentage of GL13 positive cells in at least 5-10 high power fields (x400) per sample was quantified. For each power field at least 100 cells were measured.

### Tissue Sectioning and Immunohistochemistry

Lungs and subcutaneous tumours were collected and fixed in 10% formalin overnight. Samples were then embedded in paraffin and cut in 3-7 µm thick sections. Slides were deparaffinised in xylene and re-hydrated through a series of graded ethanol until water. Antigen retrieval and immunohistochemistry was performed by the Spanish National Cancer Research Centre (CNIO) Histopathology Service. Whole slides were digitalised using an AxioScan Microscope Slide Scanner (Zeiss) using polarised light, and ZEN Blue Software (v2.6.) was used for image acquisition. Full list of antibodies used for IHC can be found in Supplementary Table 4.

### Statistics

Prism Software (GraphPad, v.9) software was used for statistical analysis. All data are displayed as mean ± SD unless stated otherwise. Group sizes were determined based on the results of preliminary experiments. Group allocation was performed in a randomized manner. Normality was tested using the Shapiro-Wilk test, and equality of variances was assessed using the F test. For normally distributed data with equal variance, statistical significance was determined using two-tailed unpaired Student’s t-test. Welch’s correction was performed for samples with unequal variance. The two-sided Mann–Whitney U-test was performed for datasets without normal distribution, one-way analysis of variance (ANOVA) was used for comparison of more than two samples and two-way ANOVA was used to analyse data with two variables, such as cell confluency or tumour growth over time. Tukey’s Multiple Comparisons test was performed to compare the mean of each group with the mean of every other group after ANOVA testing. A two-sided *log*-rank test was performed to analyse survival for categorical variables. A p-value below 0.05 was considered significant and indicated with an asterisk (*p < 0.05, **p < 0.01, ***p < 0.005, ****p < 0.001).

### Data availability

The bulk RNAseq dataset generated to support the findings of this study are available in the EMBL’s European Bioinformatics Institute (EBI) database: European Nucleotide Archive (ENA) repository, Accession Number PRJEB52271 (https://www.ebi.ac.uk/ena/browser/home). A549 MEMA data is available on Synapse (https://www.synapse.org) (ID: syn27665118). All other data supporting the findings of this study are available from the corresponding author upon request.

### Code availability

The bulk RNAseq data were aligned using STAR (v2.6.1d)^59^. Quality control was carried out with STAR (v2.6.1d) ^59^ and samtools (v1.9)^60, 61^. Read counts extracted with FeatureCounts (v1.6.4)^62^. Normalisation of read counts for analysis was carried out by EdgeR^63–65^ and Limma^66^ within R environment (v3.0.6). Pathway analyses was performed in KEGG (KEGGREST version 1.24.1)^67^, GO (GOSEQ version 1.36.0)^68^ and GSEA (FGSEA version 1.10.1)^69–71^ R packages. The A549 MEMA custom code is available on GitHub (https://github.com/markdane/A549_low_serum). RNA-seq raw data are available in the EBI database. European Nucleotide Archive (ENA) repository Accession Number PRJEB52271, and code used for data analyses are available upon request. Any additional information required to reanalyse the data reported in this work paper is available from the Lead contact upon request.

## Acknowledgements

The Muñoz-Espín laboratory is supported by the Cancer Research UK (CRUK) Cambridge Centre Early Detection Programme (RG86786), by a CRUK Programme Foundation Award (C62187/A29760), by a CRUK Early Detection OHSU Project Award (C62187/A26989), and by a Medical Research Council (MRC) New Investigator Research Grant (NIRG) (MR/R000530/1). E.G.-G. was holder of a “La Caixa” Foundation Scholarship for postgraduate studies at European Universities and funded by the CRUK Cambridge Centre Early Detection Programme. D.M. was funded by a New Investigator Research Grant (NIRG) (MR/R000530/1) and a CRUK Programme Foundation Award (C62187/A29760). S.M. is funded by a CRUK Programme Foundation Award (C62187/A29760). H-L.O. is funded by a CRUK Early Detection OHSU Project Award (C62187/A26989) and M.D. is funded by an Evelyn Trust Fellowship (G113971). V.G. is financially supported by the European Regional Development Fund of the European Union and Greek national funds through the Operational Program Competitiveness, Entrepreneurship and Innovation, under the call RESEARCH - CREATE – INNOVATE (project code: T1EDK02939) and NKUA-SARG grant 70/3/8916. This research was supported by the NIHR Cambridge Biomedical Research Centre (BRC- 1215-20014) through funding to R.C.R. and D.M.R. The views expressed are those of the authors and not necessarily those of the NIHR or the Department of Health and Social Care. G.J.D. was supported by funding through the Cancer Research UK Cambridge Centre, Thoracic Cancer Programme. M.N. and I.O. were supported by Cancer Research UK Cambridge Institute’s Core Grant (C9545/A29580) (MN) and Cancer Research UK Pioneer Award (C63389/A30462). We thank Dr David Kirsch (Duke University, UK) for the provision of the p53*^frt/frt^* mouse allele. We also thank Dr Manuel Serrano (Institute for Research in Biomedicine, Barcelona, Spain) for critical reading and insightful comments.

## Author Contributions

Project supervision and Funding acquisition: D.M-E. Conceptualisation and Design of the project: D.M.-E., E.G.-G. Experimental Designs: D.M.-E., E.G.-G, D.M. Methodology: E.G.-G., D.M., S.M. (in vivo experiments); E.G.-G., (proliferation, colony-formation, wound healing, sphere-formation, and co-culture assays; cell mitochondrial stress test; molecular biology and mechanistic assays); J.E.M. (RNAseq analyses); H-L.O. (MEMA analyses and experiments with carboplatin and PEO4 cells); M.D. (RT-qPCR experiments); I.O. (bioinformatic analyses); R.H., M.D., J.E.K. (MEMA analyses); G.M., F.M. (CT scan analyses and 3D rendering); C.P.M. (generated KP reagents and provided guidance with their *in vivo* transplantation); M.B. (oversight and generation of Kras-^+/FSFG12V^-driven lung cancer murine model); V.G. (SenTraGor analysis of human tissue sections); D.R., G.J.D., R.C.R. (provided human samples from Royal Papworth Hospital Tissue Bank and clinical oversight); M.N. (OMICS oversight and resources). Data analysis and Curation: D.M.-E., E.G.-G. Writing of the Manuscript: D.M.-E., E.G.-G. with input from all of the authors. All the authors commented on the text.

---

**Extended Data Figure 1.**
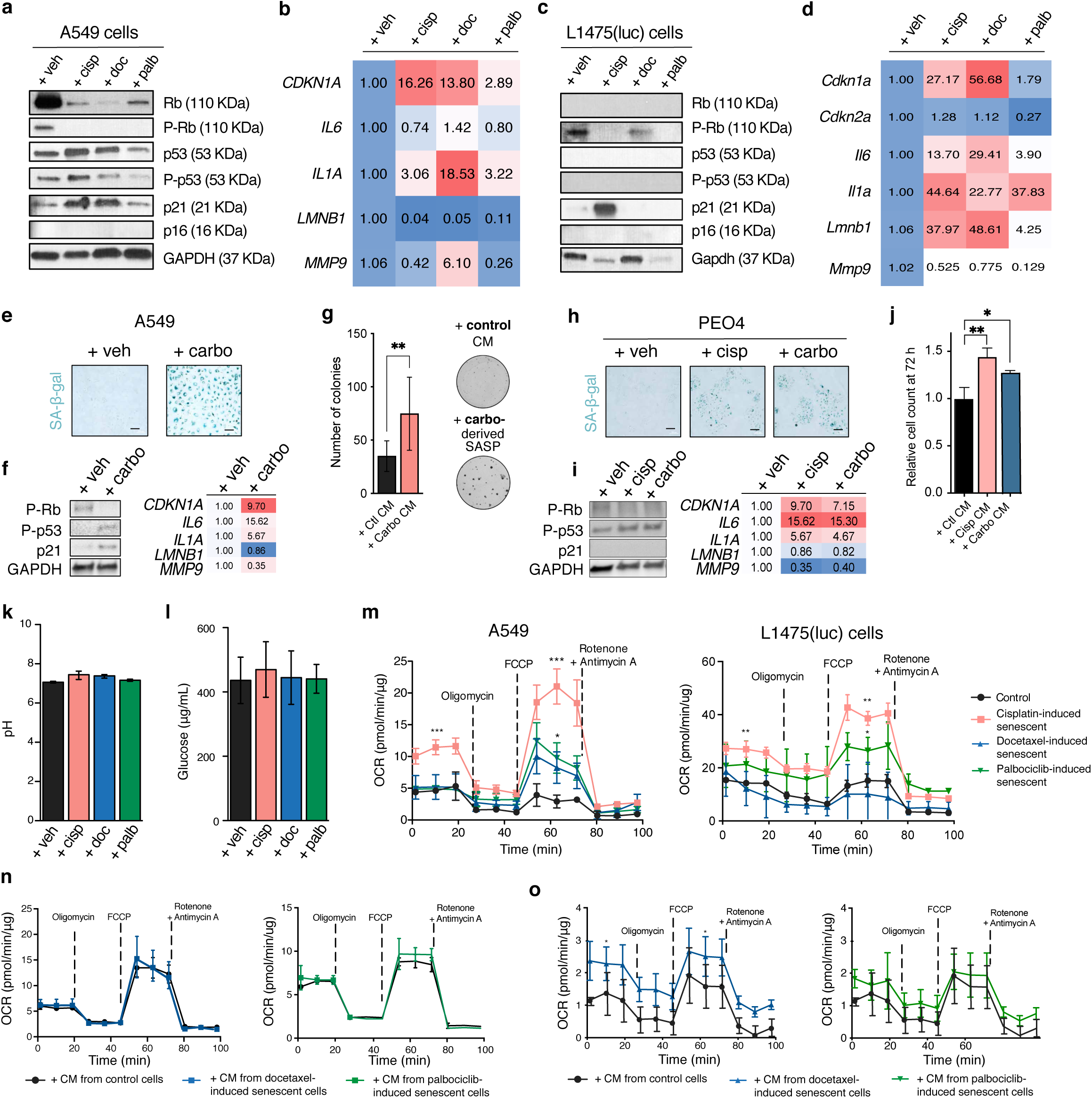
Platinum-induced senescence promotes malignant traits on non-senescent lung and ovarian cancer cells through the secretion of SASP factors. **a.** Western blot analysis for the expression of relevant senescence markers in control and senescent A549 cells upon 7 days of treatment. **b.** Fold change gene expression levels of different senescence markers in senescent A549 cells relative to vehicle-treated cells. **c**. Western blot analysis for the expression of relevant senescence markers in control and senescent L1475(luc) cells upon 5 days of treatment**. d.** Fold change gene expression levels of different senescence markers in senescent L1475(luc) cells relative to vehicle-treated cells. **e**. Representative images of control and chemotherapy-treated A549 cells fixed and stained for SA-β-gal activity after 7 days of treatment with carboplatin. Scale bar = 100 µm. **f**. Western blot analysis for the expression of relevant senescence markers in control and carboplatin-induced senescent A549 cells upon 7 days of treatment**. g**. Left, number of colonies formed upon 10 days of exposure of A549 cells to control- or carboplatin-induced senescent CM. Right, representative images of colonies for each condition. **h**. Representative images of control and chemotherapy-treated PEO4 cells fixed and stained for SA-β-gal activity after 7 days of treatment with cisplatin or carboplatin. Scale bar = 100 µm. **i**. Left, Western blot analysis for the expression of relevant senescence markers in control, cisplatin- or carboplatin-induced senescent PEO4 cells upon 7 days of treatment. Right, fold change gene expression levels of different senescence markers in senescent PEO4 cells relative to vehicle-treated cells. **j**. Relative cell count of PEO4 cells exposed to control- or chemotherapy-derived-SASPs upon 72 h exposure. **k.** Average pH measurements of the conditioned media (CM) collected from control and senescent A549 conditioned for 48 h post-removal of drug. **l.** Glucose levels of CM from control and senescent A549 cells conditioned for 48 h pot-removal of drug. **m.** Oxygen-consumption rate (OCR) of control and cisplatin-, docetaxel- and Palbociclib-induced senescent A549 (left) and L1475(luc) cells (right) in basal conditions and upon injection of oligomycin, FCCP, rotenone and antimycin A. **n.** OCR of A549 cells exposed to control and docetaxel-induced senescent cell CM (left) and control and Palbociclib-induced senescent cell CM (right) in basal conditions and upon the injection of mitochondria-targeting drugs. **o**. OCR of L1475(luc) cells exposed to control and docetaxel-induced senescent cell CM (left) and control and Palbociclib-induced senescent cell CM (right) in basal conditions and upon the injection of mitochondria-targeting drugs. Data in **a-l** are shown as mean ± SD (n = 3), and data in **m-o** are shown as mean ± SD from one representative experiment (out of three independent experiments performed). Statistical significance was assessed by two-sided, one-way or two-way ANOVA, followed by Tukey’s multiple comparisons test, *p < 0.05, **p < 0.01, ***p < 0.005.

**Extended Data Figure 2.**
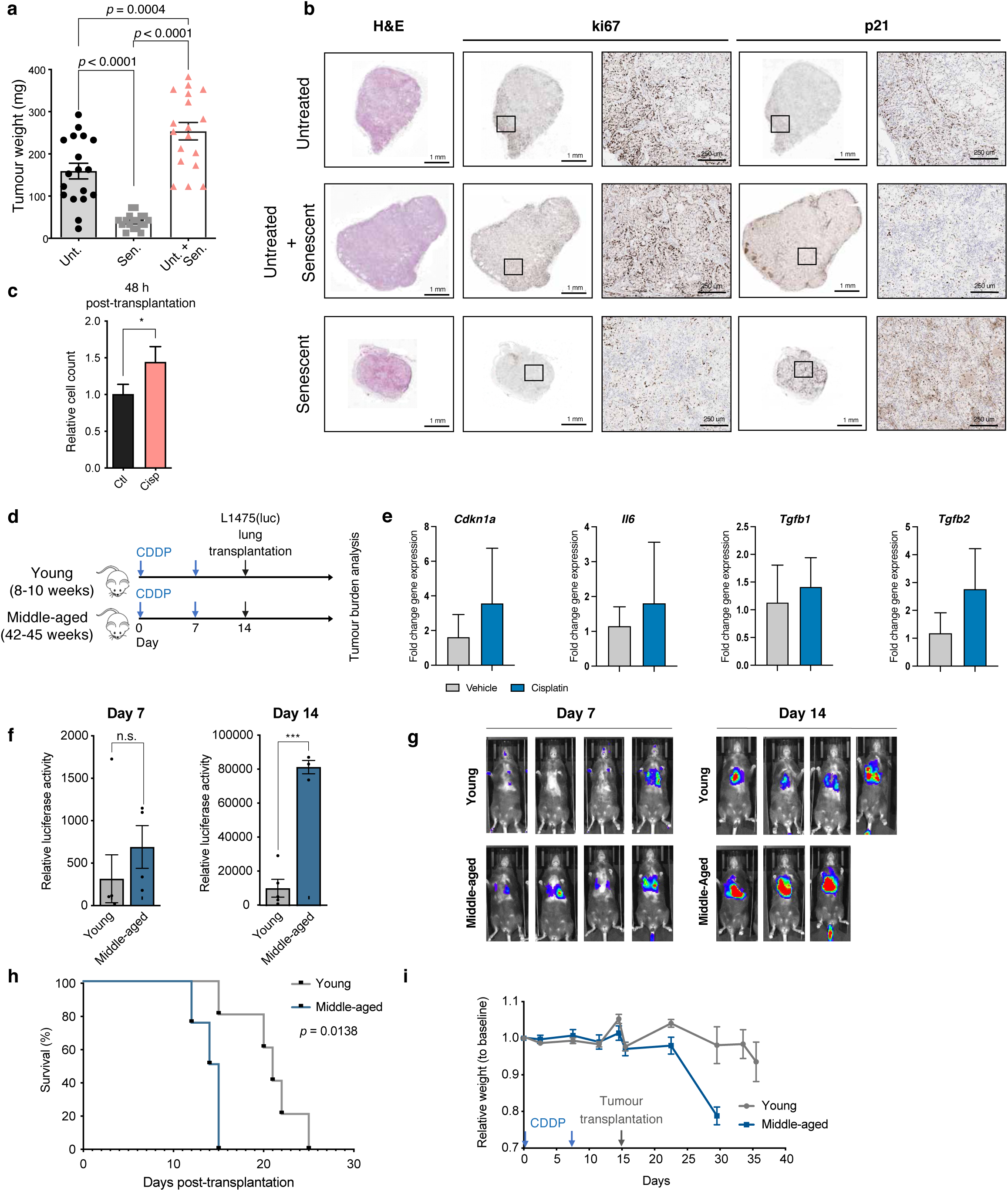
Cisplatin-induced senescence results in lung tumour promoting effects that are exacerbated during ageing. **a.** Mean tumour weight at experimental endpoint resected from mice transplanted with untreated A549 cells, a combination of untreated and cisplatin-induced senescent A549 cells or senescent-cells only (n=18 tumours per group). **b.** Representative histological images of resected xenograft from each experimental group, stained for H&E, Ki67 and p21 expression. **c.** Relative cell count of L1475(luc) cells exposed to control or cisplatin-induced senescent CM for 10 days. Assay was performed in parallel to orthotopic transplantation and cells were grown in normal media (not CM) for 48 h before cell quantification by flow cytometry. **d.** Schematic representation of experimental layout. Young (8-10-week old) and aged (42-45-week old) animals were subjected to two cycles of 1.5 mg/kg body weight CDDP treatment. Mice were then transplanted with L1475(luc) in the lungs via tail-vein injection; tumour burden was assessed twice a week by bioluminescence imaging and survival was determined as the time from tumour transplantation until the onset of moderate signs of disease. **e.** Fold change gene expression of *Cdkn1a*, *Il6*, *Tgfb1* and *Tgfb2* in samples from each experimental condition (n = 3). **f.** Quantification of luciferase activity at day 7 and 14 post-transplantation, relative to activity recorded on day 1 after cell transplantation (n = 4 per group). **g.** Representative images of luciferase activity in mice 7 and 14 days after transplantation with lung cancer cells. **h.** Survival curve of mice in the in each experimental group. **i**. Weight relative to baseline (before first CDDP dose) in each experimental group over time. Data are shown as mean ± SEM (**a, f**) and as mean ± SD (**c, i**). Statistical significance for tumour weight was assessed using one-way ANOVA followed by Tukey’s multiple comparisons test was used to assess statistical significance for tumour weight and relative weight over time in mice. Two-tailed Student’s t test was applied to assess significance for relative cell count and relative luciferase activity. Survival analysis was performed using the Kaplan-Meier method and a two-sided log-rank test was conducted to determine statistical significance. *p < 0.05, **p < 0.01, ***p < 0.005, n.s. = not significant.

**Extended Data Figure 3.**
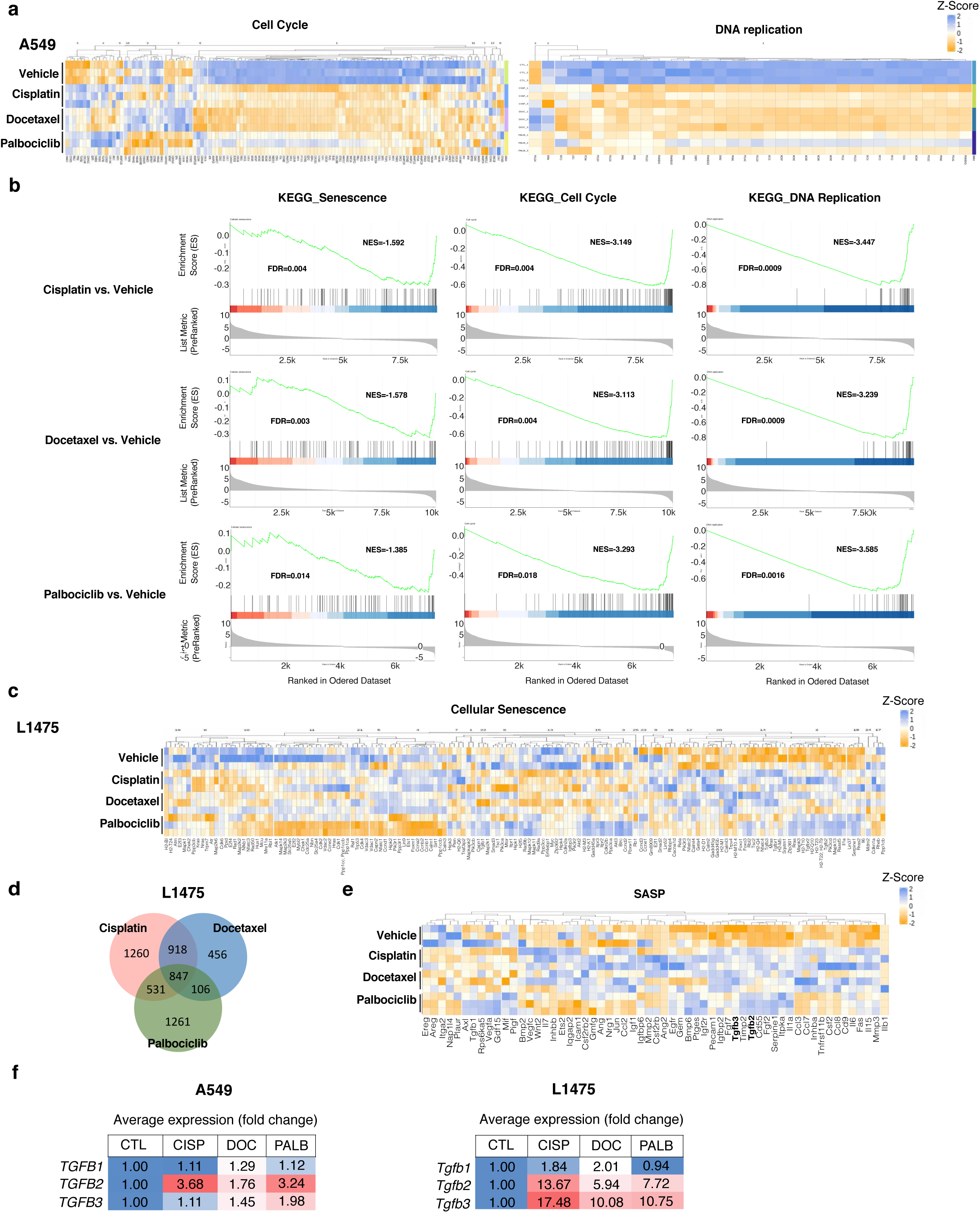
Transcriptomic profiles of A549 and L1475(luc) cells upon distinct chemotherapy treatments. **a.** Heatmap displaying expression z-scores of most significantly altered gene expression changes and their hierarchical clustering in cell cycle (left) and DNA replication pathways (right) of control and cisplatin-, docetaxel- and palbociclib-induced senescent L1475(luc) cells. **b.** GSEA of KEGG Senescence, KEGG Cell Cycle and KEGG DNA replication sets in cisplatin-, docetaxel- and palbociclib-induced senescent A549 cells versus untreated A549 cells. **c.** Heatmap displaying expression z-scores of most significantly altered gene expression changes and their hierarchical clustering in the cellular senescence pathway (mmu04218) of control and cisplatin-, docetaxel- and palbociclib-induced senescent L1475(luc) cells. **d.** Venn diagram of the number of genes significantly upregulated in cisplatin-, docetaxel- and palbociclib-induced senescent L1475(luc) cells vs control. **e.** Heatmap of expression z-scores of selected SASP genes in control and cisplatin-, docetaxel- and palbociclib-induced senescent L1475(luc) cells. **f.** Fold change TGFB1/Tgfb1, TGFB2/Tgfb2 and TGFB3/Tgfb3 gene expression levels in control and chemotherapy-treated A549 and L1475(luc) cells.

**Extended Data Figure 4.**
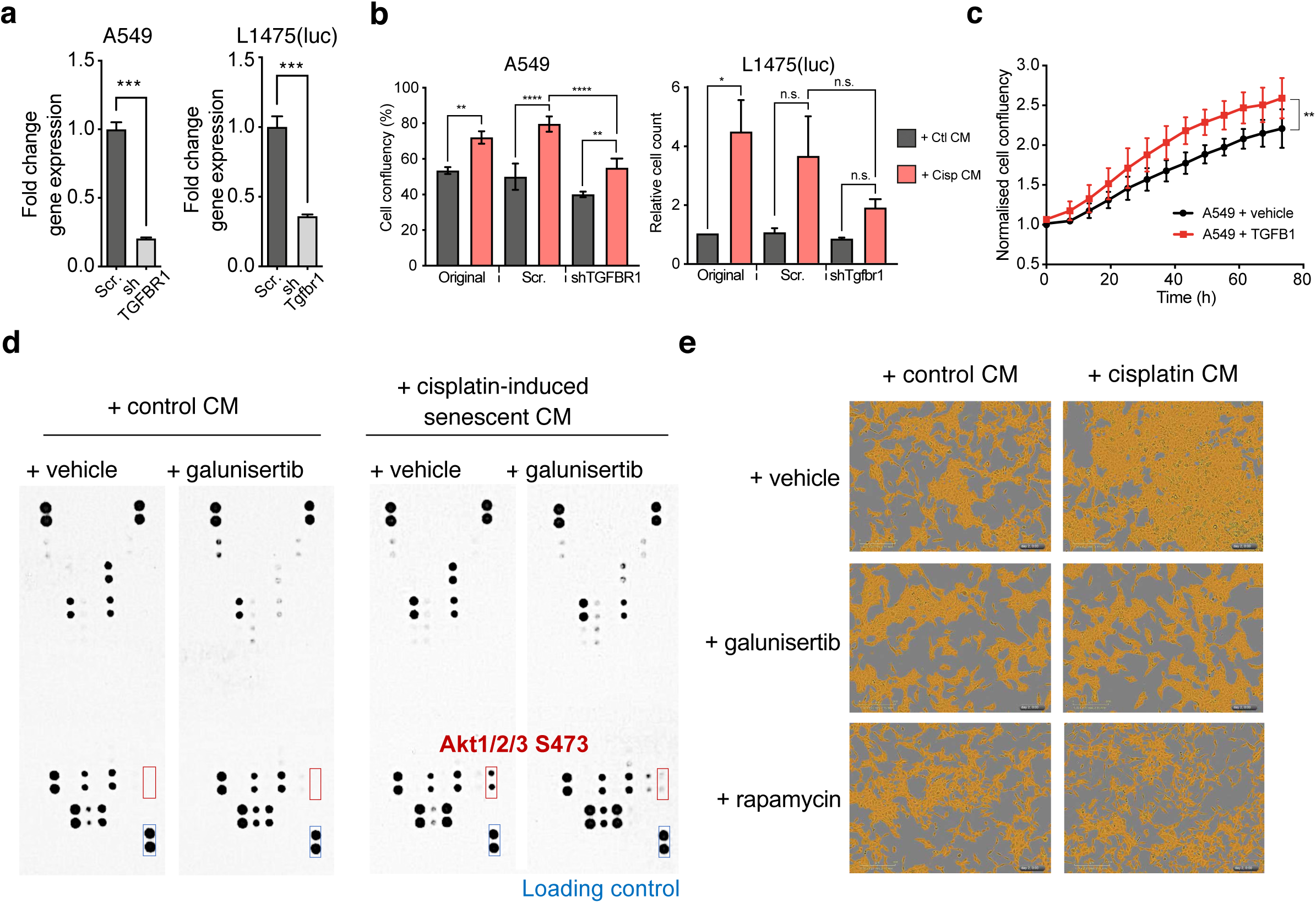
TGFβR1-driven activation of Akt/mTOR pathway orchestrates the induction of malignant traits upon exposure to cisplatin-derived SASP. **a.** Fold change TGFβR1/Tgfbr1 gene expression of scrambled- and shTGFβR1-A549 cells (left) and scrambled- and shTgfbr1-L1475(luc) cells (right). **b.** Cell confluency and relative cell count of A549 (left) and L1475(luc) (right) original, scrambled and shTGFB1/shTgfb1 cells upon exposure to control- and cisplatin-induced senescent CM for 48 h**. c.** Normalised cell confluency over time of A549 cells treated with human recombinant TGFβ1 ligand. Data in panels a-c represent mean ± SD (n=3). Statistical significance was determined by two-tailed, one- or two-way ANOVA followed by Tukey’s multiple comparisons test, and by two-tailed Student’s t-test. *p < 0.05, **p < 0.01, ***p < 0.005, n.s. = not significant. **d.** Representative pictures of human phospho-kinase array panels of whole-protein extracts from A549 cells exposed for 30 min to either control- or cisplatin-induced senescent CM with and without 50 µM galunisertib treatment. **e.** Representative images depicting A549 cell confluency from cells exposed to control- or cisplatin-induced senescent CM for 48 h treated with either vehicle, 50 µM galunisertib or 1 µM rapamycin.

**Extended Data Figure 5.**
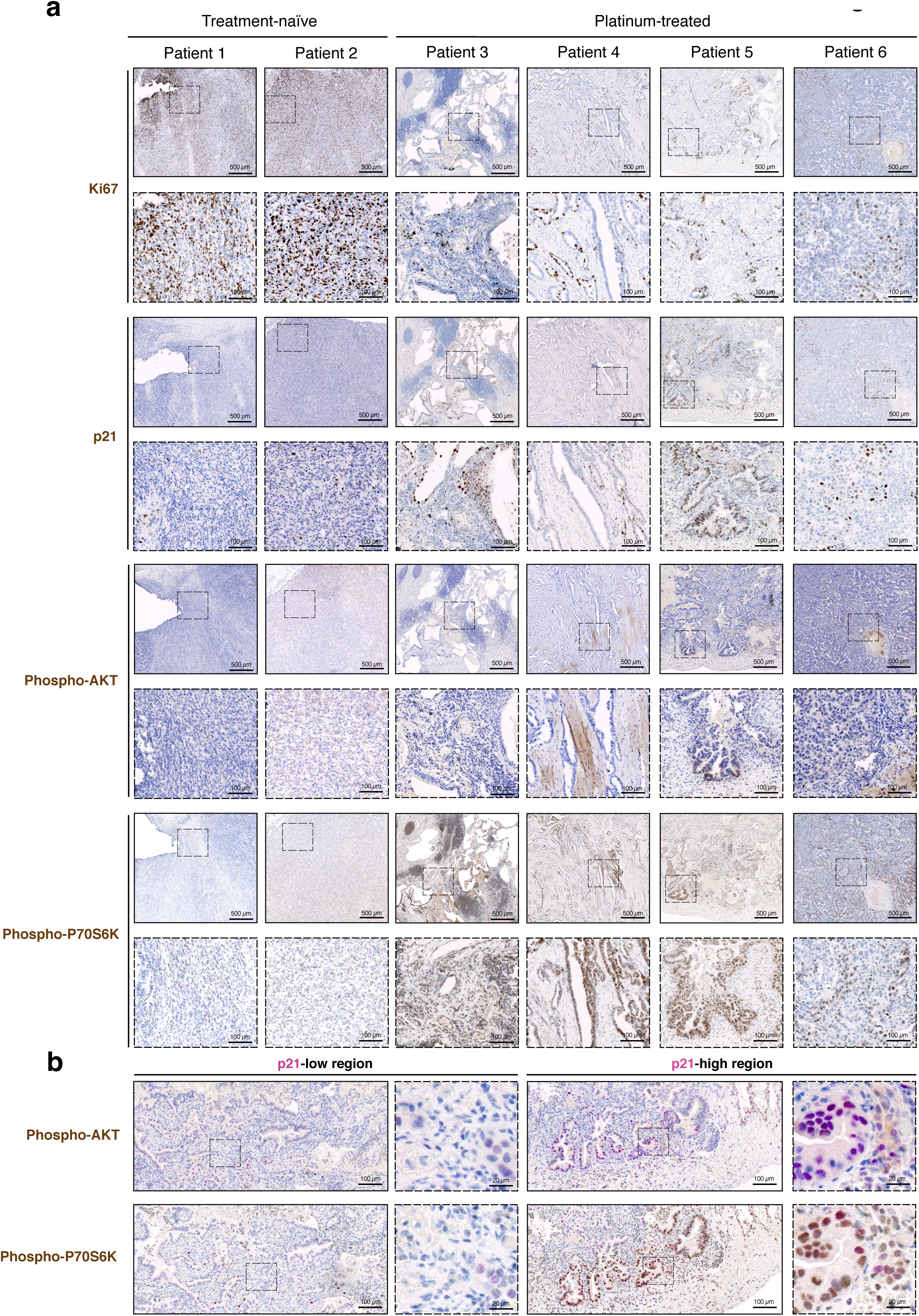
Neoadjuvant platinum-based therapy induces senescence in NSCLC patients and this correlates with increased phospho-AKT and phospho-P70S6K signalling in surrounding cells. **a.** Representative histological images of NSCLC specimens resected from treatment-naïve or platinum-treated patients and subjected to Ki67, p21, phospho-AKT and phospho- P70S6K staining. Scale bar = 500 µm or 100 µm as depicted. **b.** Representative histological images of NSCLC specimens resected from platinum-treated patients and subjected to co-staining for p21 (pink) and phospho-AKT or phospho-P70S6K (brown). Scale bar = 100 µm or 20 µm as depicted in the image.

**Extended Data Figure 6.**
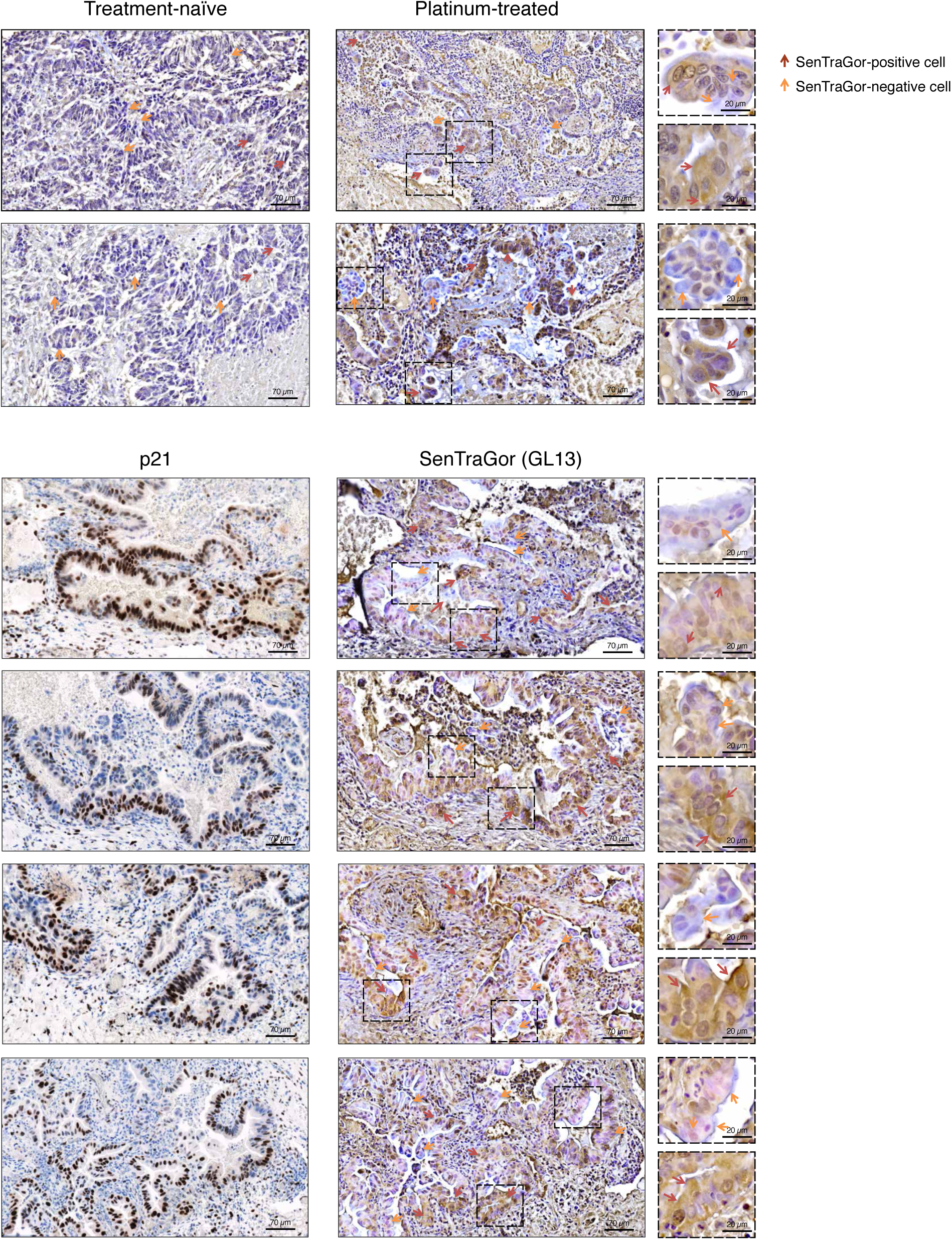
Neoadjuvant platinum-based therapy induces biomarkers of senescence in NSCLC patients. Representative histological images of NSCLC specimens resected from treatment-naïve or platinum-treated patients and subjected to SenTraGor and p21 staining. Scale bar = 70 µm or 20 µm as depicted in the image.

**Extended Data Figure 7.**
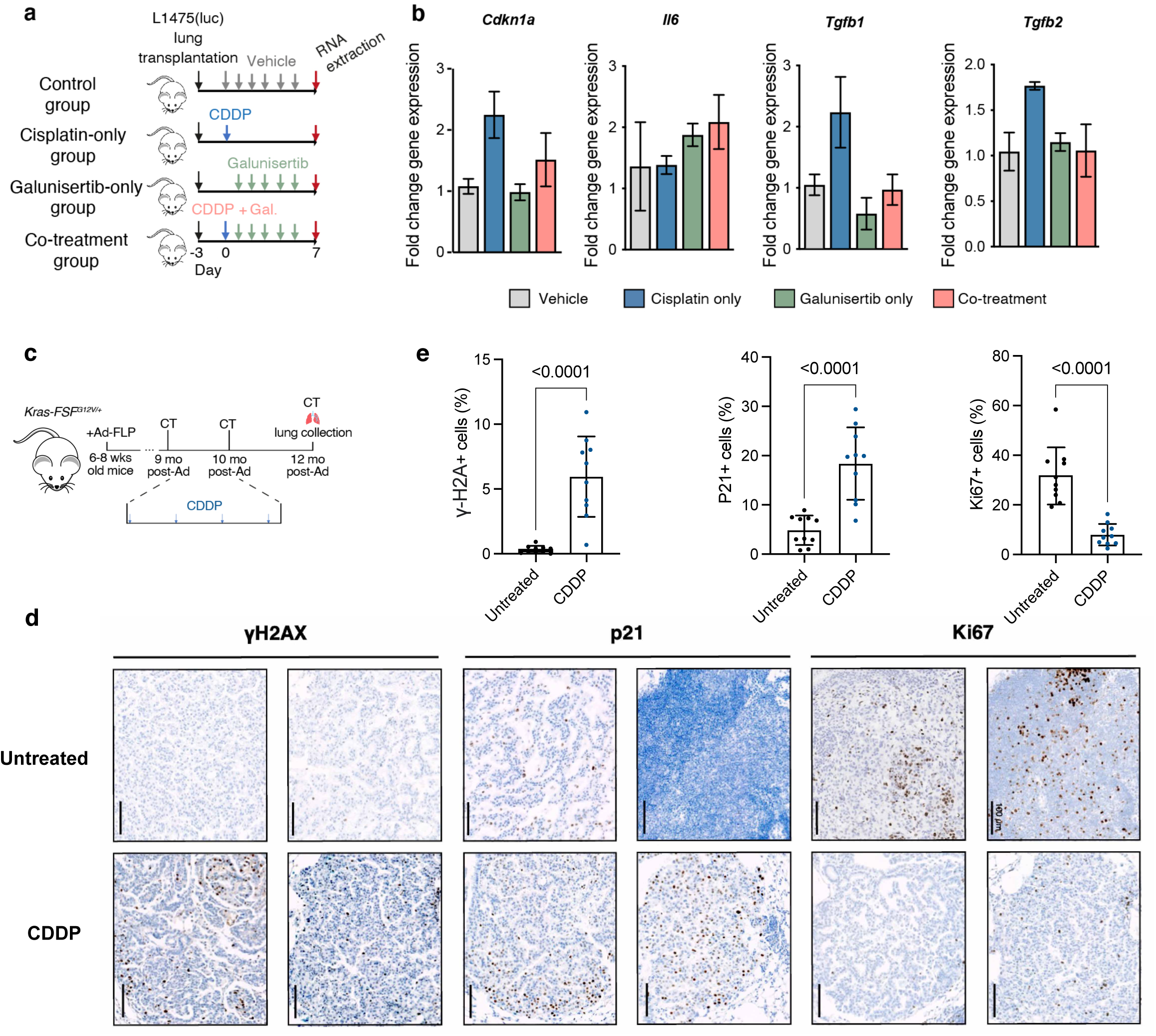
Cisplatin treatment induces senescence in Kras-driven lung cancer mouse models while galunisertib concomitant treatment reduces gene expression of senescence biomarkers and Tgfb ligands. **a.** Animals were transplanted with L1475(luc) cells in the lung via tail-vein injection, and after 3 days, they were subjected to 1.5 mg/kg body CDDP, 150 mg/kg body weight, a combination of both or vehicle as shown in the timeline. Lungs were resected at day 7 post-transplantation and RNA was extracted from whole lungs. **b.** Fold change gene expression of *Cdkn1a*, *Il6*, *Tgfb1* and *Tgfb2* in samples from each experimental condition (n = 3). Statistical significance was assessed by one-way ANOVA followed by Tukey’s multiple comparisons test. No statistically significant changes were detected. **c.** Schematic representation of experimental layout. Briefly, lung tumours were induced in 6-8 week-old Kras-FSF^G12V^ mice by Ad-FLP intratracheal injection. At 9 months post-lung cancer initiation animals were subjected to 1.5 mg/kg body CDDP once a week for 1 month. At 4 weeks upon treatment completion, animals were subjected to CT scanning for lung tumour burden assessment and histological analyses. **d.** Representative histological images of KrasFSF^G12V/+^ murine lung adenocarcinomas stained for Ki67, p21 and Hγ2AX, from animals subjected to either two doses of CDDP (1.5 mg/kg body weight, i.p.) or vehicle 9 months post-induction of tumours. **e.** Quantification of Ki67+, p21+ and Hγ2AX+ cells per total cells in treatment-naïve vehicle- or chemotherapy-treated lung tumours. For quantification, 3 representative images were quantified per individual (n=3 mice per group). Statistical significance was determined by two-tailed Student’s t-test.

**Supplementary Table 1a.** Human samples from Royal Papworth Hospital Resarch Tissue Bank.

| Patient number | TB Number | Pathological phenotype | Stage | Location | Neoadjuvant treatment |
| --- | --- | --- | --- | --- | --- |
| 1 | TB21.0150 | Lung adenocarcinoma (multifocal) | T2a N2 M0 | Right upper lobe | None |
| 2 | TB21.0151 | Lung adenocarcinoma (multifocal) | T2a N2 M0 | Left upper lobe | None |
| 3 | TB21.0210 | Lung adenocarcinoma (poorly differentiated) | T2a N2 M0 | Left upper lobe | 3 cycles of platinum-based chemotherapy |
| 4 | TB21.0209 | Lung adenocarcinoma (acinar (70%) and micropapillary (30%) growth patterns) | T3 N0 | Left upper lobe | 4 cycles of cisplatin/vinorelbine |
| 5 | TB21.0208 | Lung adenocarcinoma (multifocal with lepidic invasive acinar growth patterns) | T3 N3 M0 | Right upper lobe | 5 cycles of carboplatin/pemetrexed |
| 6 | TB14.0993 | Lung adenocarcinoma (30% lepidic component) | T3 N2 | Left upper lobe | Platinum-based chemotherapy |

**Supplementary Table 1b.**
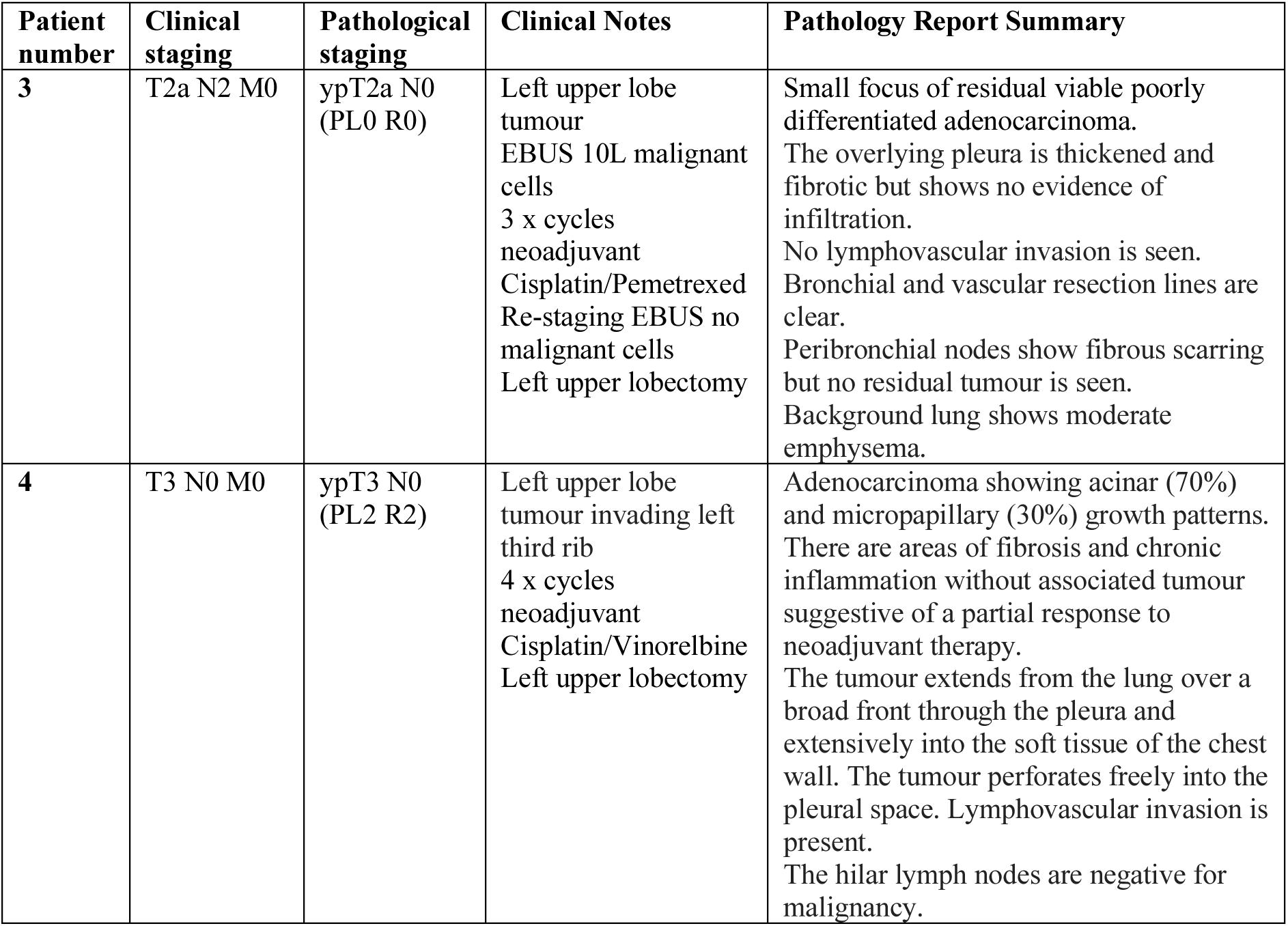
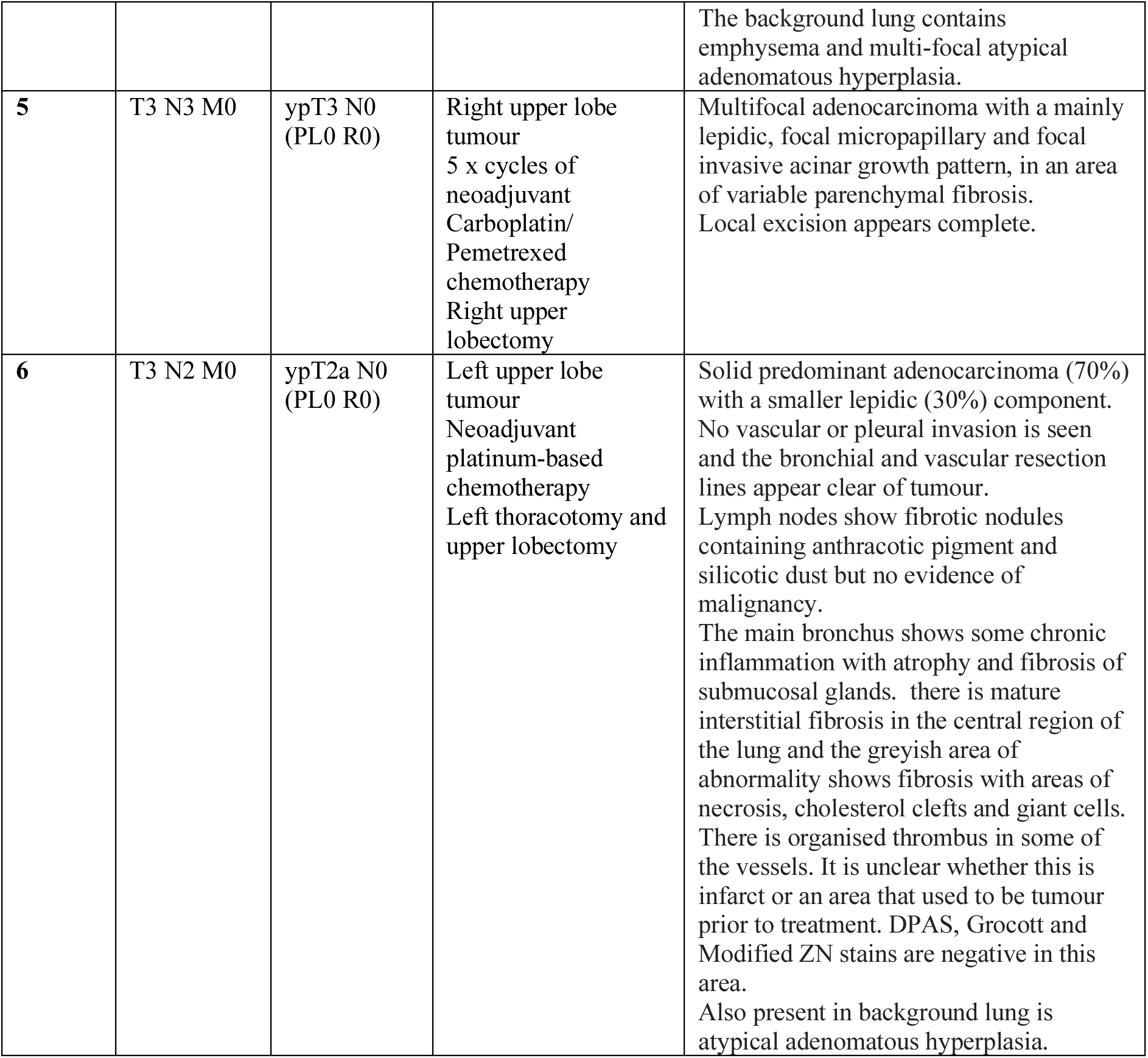
Additional clinical and pathological information for cases treated with neoadjuvant platinum-based therapy.

**Supplementary Table 2.**
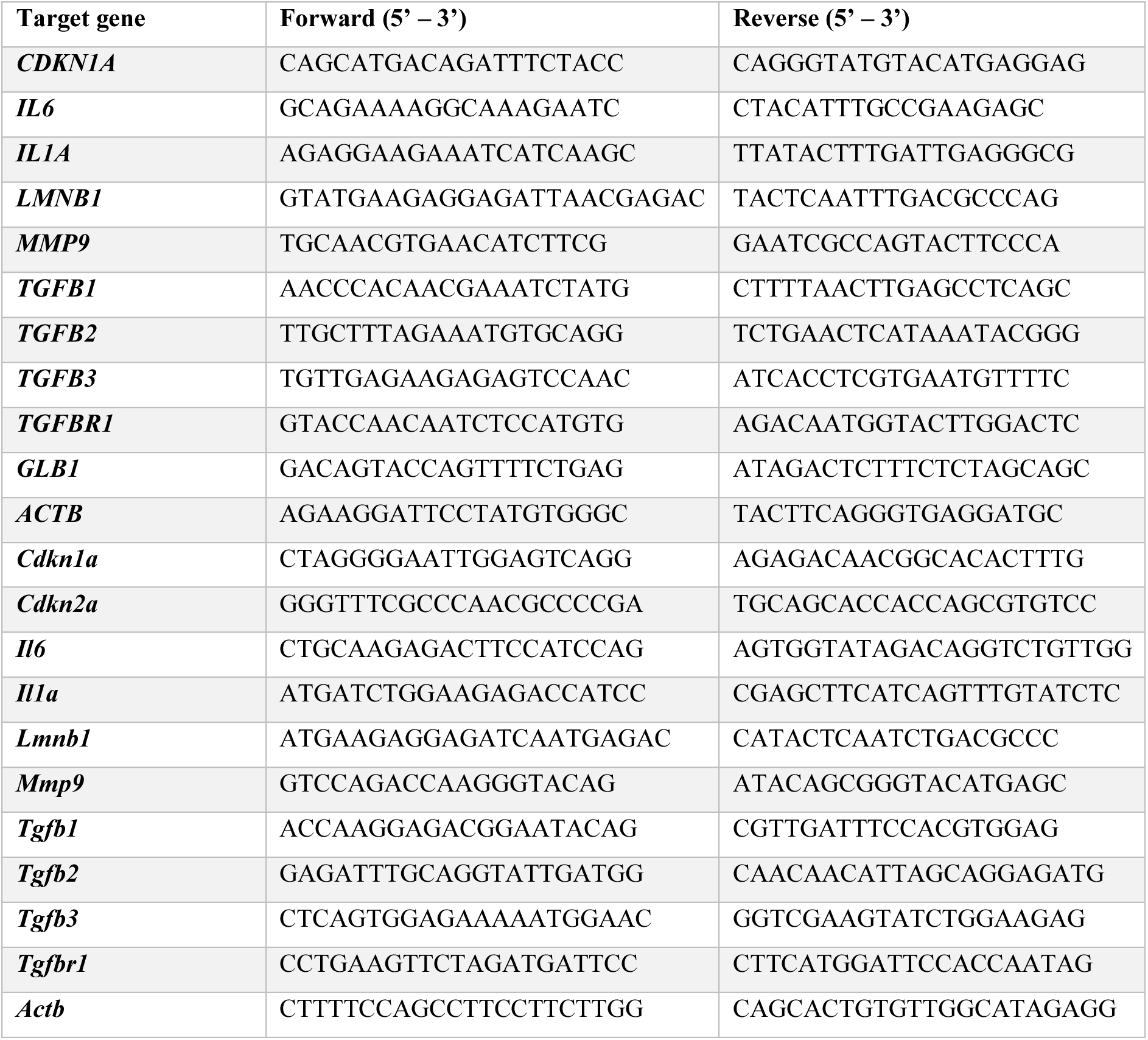
Sequences of oligonucleotides used for the amplification of target genes during RT-qPCR.

**Supplementary Table 3.**
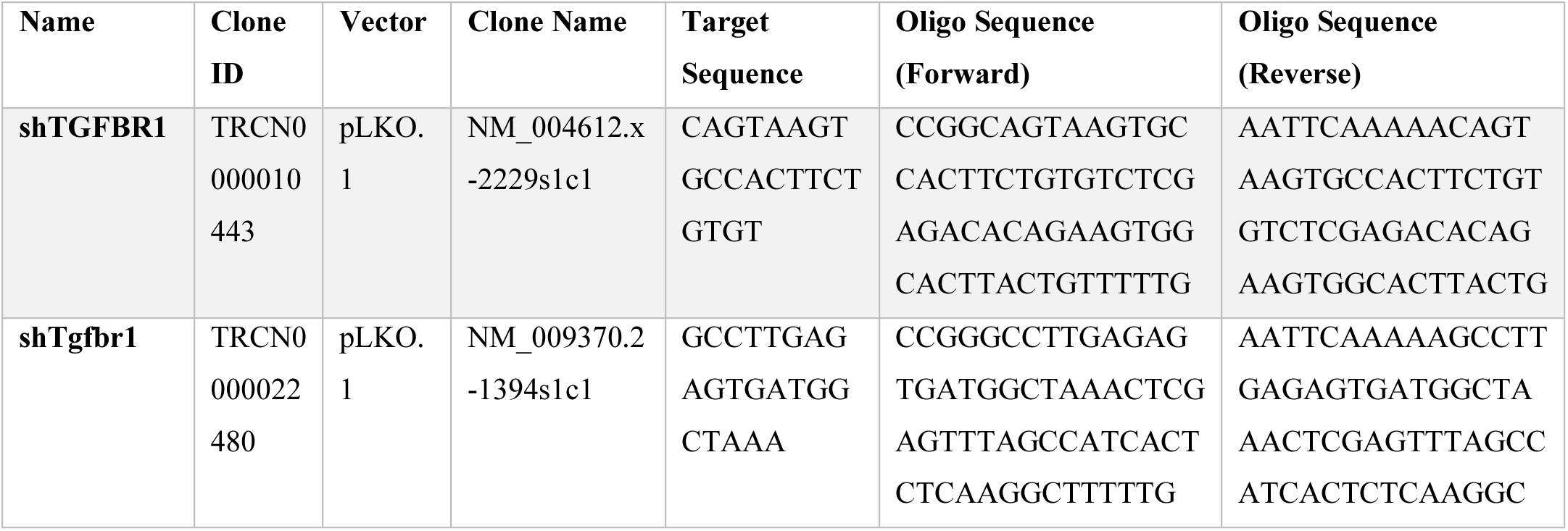
shRNA construct information and oligo sequences.

**Supplementary Table 4.** List of primary antibodies used for Western Blotting.

| Antibody | Species | Provider | Catalogue Number | Working Dilution |
| --- | --- | --- | --- | --- |
| Anti-GAPDH | Rabbit | Abcam | ab9485 | 1:2000 |
| Anti-Rb (4H1) | Mouse | Cell Signaling | 9309 | 1:2000 |
| Anti-Phospho-Rb (Ser807/811) | Rabbit | Cell Signaling | 9308 | 1:1000 |
| Anti-p53 (CO-7) | Mouse | Santa Cruz<br>Biotechnology | sc-47698 | 1:500 |
| Anti-Phospho-p53 (Ser20) | Rabbit | Cell Signaling | 9287 | 1:1000 |
| Anti-p21 (F-5) | Goat | Santa Cruz<br>Biotechnology | sc-6246 | 1:250 |
| Anti-CDKN2A/p16INK4a (EPR1473) | Rabbit | Abcam | ab108349 | 1:2000 |
| Anti-TGFB1 | Rabbit | Abcam | ab92486 | 1:1000 |
| Anti-TGFB2 | Mouse | Abcam | ab36495 | 1:1000 |
| Anti-TGFB3 | Goat | Bio-Techne | AF-243-NA | 1:1000 |
| Anti-Pan-P70S6K | Mouse | Bio-Techne | MAB8962 | 1:1000 |
| Anti-Phospho-p70S6K (T389) | Rabbit | Bio-Techne | MAB8963 | 1:1000 |
| Anti-Pan-Akt | Rabbit | Cell Signaling | 4691 | 1:1000 |
| Anti-Phospho-Akt (S473) | Rabbit | Cell Signaling | 9271 | 1:1000 |
| Anti-TGFBR1 | Goat | Abcam | ab121024 | 1:1000 |

List of secondary antibodies used for Western Blotting.
| Antibody | Species | Provider | Catalogue Number | Working Dilution |
| --- | --- | --- | --- | --- |
| HRP-conjugated AffiniPute Anti-Rabbit IgG (H+L) | Donkey | Jackson<br>ImmunoResearch | 711-035-152 | 1:5000 |
| HRP-conjugated AffiniPute Anti-Mouse IgG (H+L) | Donkey | Jackson<br>ImmunoResearch | 715-035-150 | 1:5000 |
| HRP-conjugated AffiniPute Anti-Goat IgG (H+L) | Donkey | Jackson<br>ImmunoResearch | 805-035-180 | 1:5000 |

List of primary antibodies used for IHC.
| Antibody | Provider | Target Species | Catalogue Number | Clonal (Clone) |
| --- | --- | --- | --- | --- |
| Ki-67 | Dako | Human | IR626 | Mouse monoclonal (MIB-1) |
| Ki-67 | Cell Signaling | Mouse | 12202 | Rabbit monoclonal (D3B5) |
| p21 | Dako | Human | M7202 | Mouse monoclonal (CO-7) |
| **p16** | Abcam | Human | ab81278 | Rabbit monoclonal (EP435Y-129R) |
| **Phospho-Histone H2A.X (Ser139)** | Millipore | Mouse | 05-636 | Mouse monoclonal (JBW301) |
| **Cleaved Caspase-3** | Cell Signaling | Mouse | 9661 | Rabbit Polyclonal (Asp175) |
| **Phospho-AKT (Ser473)** | Cell Signaling | Human | 4060 | Rabbit Monoclonal (D9E) |
| **Phospho-p70 S6 Kinase** | Cell Signaling | Human | 9206 | Mouse Monoclonal (1A5) |
| **Biotin** | Abcam | Human | ab234284 | Rabbit Monoclonal (BTN/2032R) |
| **KRT5** | Biolegend | Human | 905051 | Rabbit Polyclonal (Poly19055) |
| **KRT19** | Millipore | Human | Mab3238 | Mouse monoclonal (RCK108) |

**Supplementary Table 5.**
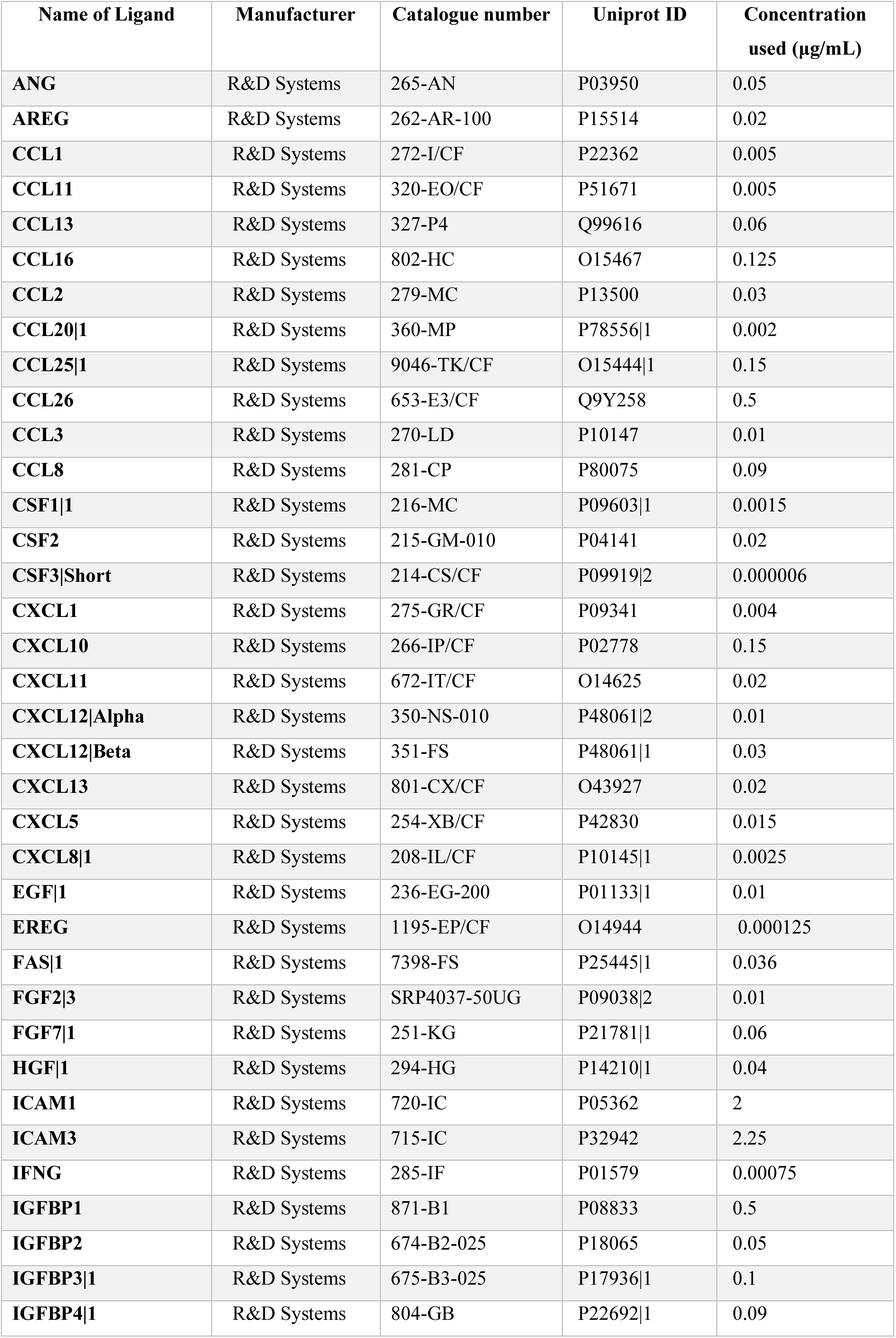
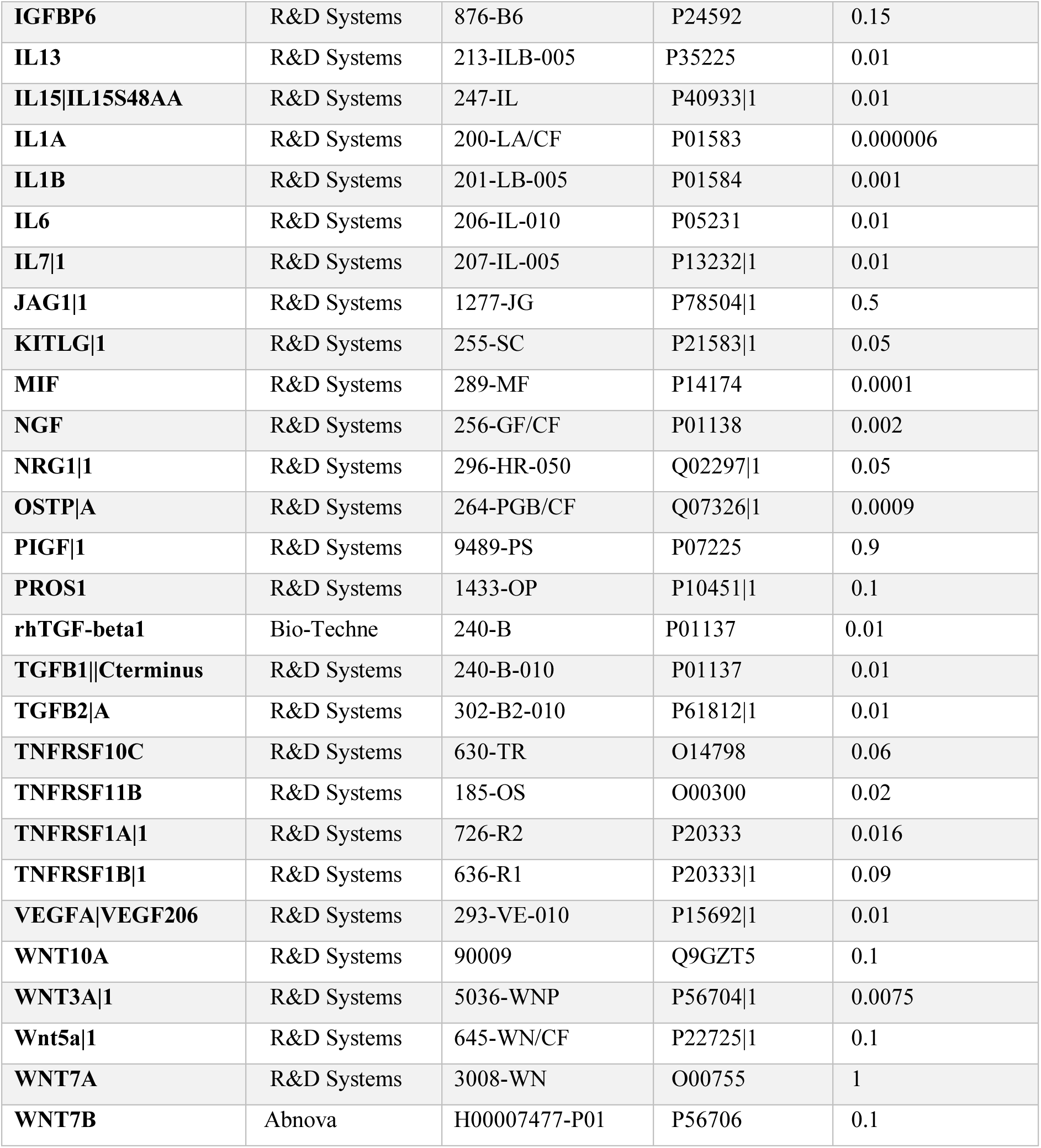
List of recombinant protein ligands used in MEMA experiments.

| Name of Ligand | Manufacturer | Catalogue number | Uniprot ID | Concentration used (µg/mL) |
| --- | --- | --- | --- | --- |
| ANG | R&D Systems | 265-AN | P03950 | 0.05 |
| AREG | R&D Systems | 262-AR-100 | P15514 | 0.02 |
| CCL1 | R&D Systems | 272-I/CF | P22362 | 0.005 |
| CCL11 | R&D Systems | 320-EO/CF | P51671 | 0.005 |
| CCL13 | R&D Systems | 327-P4 | Q99616 | 0.06 |
| CCL16 | R&D Systems | 802-HC | O15467 | 0.125 |
| CCL2 | R&D Systems | 279-MC | P13500 | 0.03 |
| CCL20\|1 | R&D Systems | 360-MP | P78556\|1 | 0.002 |
| CCL25\|1 | R&D Systems | 9046-TK/CF | O15444\|1 | 0.15 |
| CCL26 | R&D Systems | 653-E3/CF | Q9Y258 | 0.5 |
| CCL3 | R&D Systems | 270-LD | P10147 | 0.01 |
| CCL8 | R&D Systems | 281-CP | P80075 | 0.09 |
| CSF1\|1 | R&D Systems | 216-MC | P09603\|1 | 0.0015 |
| CSF2 | R&D Systems | 215-GM-010 | P04141 | 0.02 |
| CSF3\|Short | R&D Systems | 214-CS/CF | P09919\|2 | 0.000006 |
| CXCL1 | R&D Systems | 275-GR/CF | P09341 | 0.004 |
| CXCL10 | R&D Systems | 266-IP/CF | P02778 | 0.15 |
| CXCL11 | R&D Systems | 672-IT/CF | O14625 | 0.02 |
| CXCL12\|Alpha | R&D Systems | 350-NS-010 | P48061\|2 | 0.01 |
| CXCL12\|Beta | R&D Systems | 351-FS | P48061\|1 | 0.03 |
| CXCL13 | R&D Systems | 801-CX/CF | O43927 | 0.02 |
| CXCL5 | R&D Systems | 254-XB/CF | P42830 | 0.015 |
| CXCL8\|1 | R&D Systems | 208-IL/CF | P10145\|1 | 0.0025 |
| EGF\|1 | R&D Systems | 236-EG-200 | P01133\|1 | 0.01 |
| EREG | R&D Systems | 1195-EP/CF | O14944 | 0.000125 |
| FAS\|1 | R&D Systems | 7398-FS | P25445\|1 | 0.036 |
| FGF2\|3 | R&D Systems | SRP4037-50UG | P09038\|2 | 0.01 |
| FGF7\|1 | R&D Systems | 251-KG | P21781\|1 | 0.06 |
| HGF\|1 | R&D Systems | 294-HG | P14210\|1 | 0.04 |
| ICAM1 | R&D Systems | 720-IC | P05362 | 2 |
| ICAM3 | R&D Systems | 715-IC | P32942 | 2.25 |
| IFNG | R&D Systems | 285-IF | P01579 | 0.00075 |
| IGFBP1 | R&D Systems | 871-B1 | P08833 | 0.5 |
| IGFBP2 | R&D Systems | 674-B2-025 | P18065 | 0.05 |
| IGFBP3\|1 | R&D Systems | 675-B3-025 | P17936\|1 | 0.1 |
| IGFBP4\|1 | R&D Systems | 804-GB | P22692\|1 | 0.09 |
| **IGFBP6** | R&D Systems | 876-B6 | P24592 | 0.15 |
| **IL13** | R&D Systems | 213-ILB-005 | P35225 | 0.01 |
| **IL15\|IL15S48AA** | R&D Systems | 247-IL | P40933\|1 | 0.01 |
| **IL1A** | R&D Systems | 200-LA/CF | P01583 | 0.000006 |
| **IL1B** | R&D Systems | 201-LB-005 | P01584 | 0.001 |
| **IL6** | R&D Systems | 206-IL-010 | P05231 | 0.01 |
| **IL7\|1** | R&D Systems | 207-IL-005 | P13232\|1 | 0.01 |
| **JAG1\|1** | R&D Systems | 1277-JG | P78504\|1 | 0.5 |
| **KITLG\|1** | R&D Systems | 255-SC | P21583\|1 | 0.05 |
| **MIF** | R&D Systems | 289-MF | P14174 | 0.0001 |
| **NGF** | R&D Systems | 256-GF/CF | P01138 | 0.002 |
| **NRG1\|1** | R&D Systems | 296-HR-050 | Q02297\|1 | 0.05 |
| **OSTP\|A** | R&D Systems | 264-PGB/CF | Q07326\|1 | 0.0009 |
| **PIGF\|1** | R&D Systems | 9489-PS | P07225 | 0.9 |
| **PROS1** | R&D Systems | 1433-OP | P10451\|1 | 0.1 |
| **rhTGF-beta1** | Bio-Techne | 240-B | P01137 | 0.01 |
| **TGFB1\|\|Cterminus** | R&D Systems | 240-B-010 | P01137 | 0.01 |
| **TGFB2\|A** | R&D Systems | 302-B2-010 | P61812\|1 | 0.01 |
| **TNFRSF10C** | R&D Systems | 630-TR | O14798 | 0.06 |
| **TNFRSF11B** | R&D Systems | 185-OS | O00300 | 0.02 |
| **TNFRSF1A\|1** | R&D Systems | 726-R2 | P20333 | 0.016 |
| **TNFRSF1B\|1** | R&D Systems | 636-R1 | P20333\|1 | 0.09 |
| **VEGFA\|VEGF206** | R&D Systems | 293-VE-010 | P15692\|1 | 0.01 |
| **WNT10A** | R&D Systems | 90009 | Q9GZT5 | 0.1 |
| **WNT3A\|1** | R&D Systems | 5036-WNP | P56704\|1 | 0.0075 |
| **Wnt5a\|1** | R&D Systems | 645-WN/CF | P22725\|1 | 0.1 |
| **WNT7A** | R&D Systems | 3008-WN | O00755 | 1 |
| **WNT7B** | Abnova | H00007477-P01 | P56706 | 0.1 |

